# MARCO^+^ lymphatic endothelial cells sequester arboviruses to limit viremia and viral dissemination

**DOI:** 10.1101/2021.06.10.447957

**Authors:** Kathryn S. Carpentier, Ryan M. Sheridan, Cormac J. Lucas, Bennett J. Davenport, Frances S. Li, Erin D. Lucas, Mary K. McCarthy, Glennys V. Reynoso, Nicolas A. May, Beth A.J. Tamburini, Jay R. Hesselberth, Heather D. Hickman, Thomas E. Morrison

## Abstract

While viremia in the vertebrate host is a major determinant of arboviral reservoir competency, transmission efficiency, and disease severity, immune mechanisms that control arboviral viremia are poorly defined. Here, we identify critical roles for the scavenger receptor MARCO in controlling viremia during arthritogenic alphavirus infections in mice. Following subcutaneous inoculation, alphavirus particles drain via the lymph and are rapidly captured by MARCO^+^ lymphatic endothelial cells (LECs) in the draining lymph node (dLN), limiting viral spread to the bloodstream. Upon reaching the bloodstream, alphavirus particles are cleared from the circulation by MARCO-expressing Kupffer cells in the liver, limiting viremia and further viral dissemination. MARCO-mediated accumulation of alphavirus particles in the dLN and liver is an important host defense mechanism as viremia and viral tissue burdens are elevated in MARCO^-/-^ mice and disease is more severe. These findings uncover a previously unrecognized arbovirus scavenging role for LECs and improve our mechanistic understanding of viremia control during arboviral infections.

## Introduction

Over the last two decades we have experienced the unanticipated emergence or re-emergence of multiple arboviruses, leading to far-reaching epidemics. In 2004, chikungunya virus (CHIKV), a mosquito-borne alphavirus, re-emerged in the Indian Ocean region and has since infected millions of people in epidemics spanning the globe, including the Americas (Moro *et al*, 2010; Volk *et al*, 2010; Zeller *et al*, 2016). CHIKV and closely related alphaviruses (e.g., Mayaro, o’nyong’nyong (ONNV), and Ross River (RRV) viruses) cause severe arthralgia and arthritis affecting the small joints. These debilitating symptoms can persist for months to years after infection (Borgherini *et al*, 2008; Couturier *et al*, 2012; Rodríguez-Morales *et al*, 2016; Schilte *et al*, 2013), and have severe economic consequences (Cardona-Ospina *et al*, 2015; Soumahoro *et al*, 2011; Vijayakumar *et al*, 2013). In 2007, the previously obscure Zika virus (ZIKV) caused multiple outbreaks in islands of the Pacific Ocean before spreading to the Americas in 2015 (Duffy *et al*, 2009; Metsky *et al*, 2017; Musso *et al*, 2018). This epidemic revealed an unexpected association of ZIKV with severe disease manifestations, including Guillain-Barré syndrome and congenital ZIKV syndrome (Pierson & Diamond, 2018). Because of this, in 2016 the WHO declared the ZIKV outbreak in the Americas a Public Health Emergency of International Concern. These events underscore the ongoing threat that zoonotic arboviruses pose.

Arboviral infections in humans are often the result of spillover from enzootic cycles, and for many arboviruses, humans are a dead-end host. However, some arboviruses sustain human-mosquito-human transmission, including dengue virus (DENV), yellow fever virus (YFV), ZIKV, and CHIKV (Weaver, 2018), which facilitates global emergence through air travel and allows for rapid spread of the virus through urban areas. While there are many determinants of arbovirus urbanization, a key factor is the development of a magnitude and duration of viremia sufficient to support infection of mosquitoes (Weaver, 2018). Beyond influencing reservoir competency and transmission efficiency, viremia also positively correlates with arboviral disease severity (Chow *et al*, 2011; de St Maurice *et al*, 2018; Pozo-Aguilar *et al*, 2014; Vaughn *et al*, 2000; Vuong *et al*, 2020; Waggoner *et al*, 2016). Thus, understanding factors that influence the magnitude and duration of viremia following arboviral infection is of critical importance.

Following the delivery of arboviruses via a mosquito bite, the virus replicates at the site of inoculation before spreading via the lymph to ultimately reach the bloodstream (Johnston *et al*, 2000; MacDonald, 2000). Given this, arboviruses must evade immune defenses in draining lymph nodes (dLNs) to establish a primary viremia. Within the dLN, subcapsular sinus (SCS) macrophages and medullary sinus (MS) macrophages are strategically positioned to encounter lymph-borne pathogens (Bellomo *et al*, 2018). These cells have been described as molecular “flypaper” given their roles in rapidly capturing a wide range of incoming particulate antigen, including lymph-borne virions (Farrell *et al*, 2015; Junt *et al*, 2007). Moreover, viral replication within SCS macrophages initiates interferon production and facilitates recruitment and activation of immune cells to limit further viral dissemination (Iannacone *et al*, 2010; Kastenmuller *et al*, 2012).

Upon reaching the bloodstream, virus particles must evade clearance by blood-filtering organs to maintain viremia and disseminate to distal tissues. The liver and spleen contain phagocytic cells strategically positioned to recognize and remove circulating self and non-self molecules. In the splenic marginal zone, marginal zone (MZM) and metallophilic (MMM) macrophages remove circulating apoptotic cells, antigen and microbes (Borges da Silva *et al*, 2015; Lewis *et al*, 2019). In the liver, Kupffer cells (KCs), which account for 80-90% of all tissue macrophages (Bilzer *et al*, 2006), line the sinusoids to detect and clear blood-borne microbes and modified host molecules from the circulation (Hickey & Kubes, 2009; Lee *et al*, 2010; Zeng *et al*, 2016).

In prior studies, we found that i.v. inoculated arthritogenic alphavirus virions were rapidly removed from the circulation and accumulated in the liver (Carpentier *et al*, 2019). In addition, we identified the scavenger receptor MARCO as essential for alphavirus particle clearance from the blood (Carpentier *et al*., 2019). These findings revealed a critical host defense mechanism that contributes to the control of arbovirus viremia once viral particles have reached the bloodstream. Here, using genetic approaches, confocal microscopy, and single cell mRNA sequencing (scRNA-seq) analysis of dLN cell populations, we expand these analyses to improve our mechanistic understanding of the role of MARCO during arthritogenic alphavirus infection following a more natural subcutaneous inoculation route. Our studies revealed two distinct roles for MARCO in controlling arthritogenic alphavirus dissemination in vertebrate hosts. First, MARCO^+^ LECs in the dLN sequester CHIKV particles to delay the establishment of viremia. Once this barrier is breached, MARCO-expressing Kupffer cells in the liver provide a second layer of protection by removing circulating viral particles. These findings advance our understanding of the immune mechanisms that control arthritogenic alphavirus viremia and dissemination and reveal an arbovirus-scavenging role for LECs.

## Results

### CHIKV infection outcomes are more severe in MARCO^-/-^ mice

To further elucidate the role of MARCO during arthritogenic alphavirus infection, we inoculated four-week-old WT or MARCO^-/-^ mice subcutaneously (s.c.) in the left rear footpad with CHIKV and evaluated disease outcomes. As a control, WT and MARCO^-/-^ mice were inoculated with CHIKV E2 K200R, a mutant virus that evades MARCO-mediated clearance from the circulation (Carpentier *et al*., 2019) and causes severe disease in WT mice (Hawman *et al*, 2017). WT mice inoculated with WT CHIKV steadily gained weight (**Fig 1A**) and displayed little to no defects in gait or hind-limb gripping ability (**Fig 1B**). In contrast, and similar to previous findings (Hawman *et al*., 2017), WT mice inoculated with CHIKV E2 K200R had delayed weight gain and developed more severe signs of musculoskeletal disease. Similarly, MARCO^-/-^ mice infected with either WT CHIKV or CHIKV E2 K200R developed more severe disease signs (**Fig 1A** and **1B**). Notably, the disease observed in MARCO^-/-^ mice infected with CHIKV E2 K200R was not more severe than disease in MARCO^-/-^ mice infected with WT CHIKV, suggesting that the enhanced disease caused by CHIKV E2 K200R in WT mice is due to evasion of MARCO. These findings demonstrate that the scavenger receptor MARCO protects from severe CHIKV disease.

**Figure 1.**
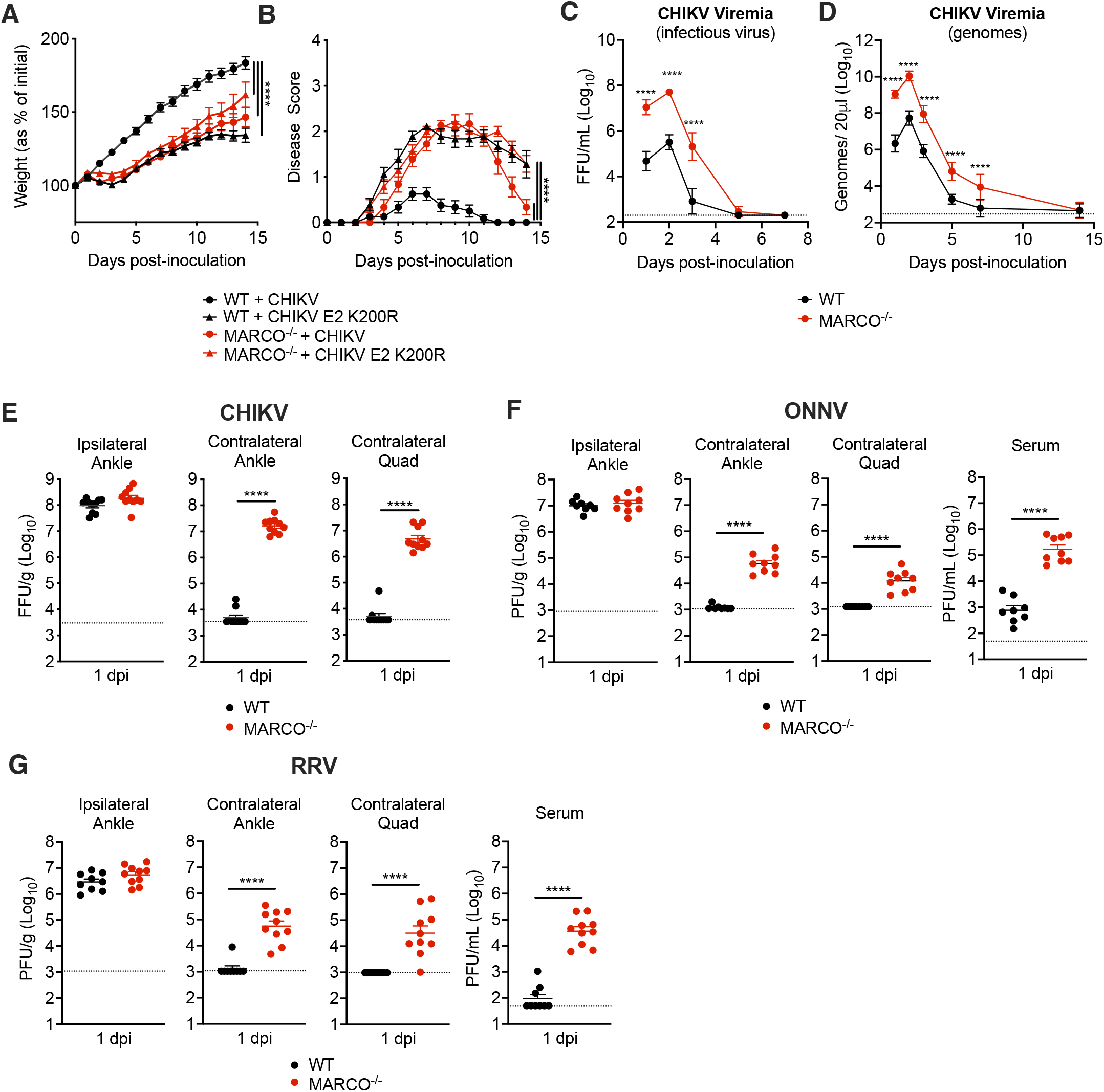
CHIKV infection outcomes are more severe in MARCO^-/-^ mice and viremia and dissemination are enhanced. **(A-B)** WT or MARCO^-/-^ C57BL/6 mice were inoculated subcutaneously (s.c.) in the left rear footpad with 10^3^ PFU of CHIKV or CHIKV E2 K200R. The percent of starting body weight **(A)** and disease score (**B**) were recorded daily over 14 days. Mean ± SEM. N=7-12, three experiments. Two-way ANOVA with Bonferroni’s multiple comparison test; *\*\*\*\*P <* 0.0001. (**C-E**) WT or MARCO^-/-^ C57BL/6 mice were inoculated s.c. in the left rear footpad with 10^3^ PFU of CHIKV. Serum collected on days 1, 2, 3, 5, 7, and 14 was analyzed by focus formation assay (FFA), mean ± SD **(C)** and by RT-qPCR, mean ± SD **(D)**. **(E)** Viral tissue burdens were analyzed at 1 day post inoculation (dpi) by FFA. Mean ± SEM. Two experiments, n=5-10. Two-way ANOVA with Bonferroni’s multiple comparison test **(C-D**) or Mann-Whitney test **(E**); \*\*\*\**P* < 0.0001. **(F-G)** WT or MARCO^-/-^ C57BL/6 mice were inoculated s.c. in the left rear footpad with 10^3^ PFU of ONNV **(F)** or RRV **(G).** Tissues and serum were collected at 1 dpi and analyzed by plaque assay. Mean ± SEM. Two experiments, n= 8-10. Mann-Whitney test; \*\*\*\**P* < 0.0001.

### CHIKV viremia and tissue burdens are elevated in MARCO^-/-^ mice

Since our previous work identified a critical role for MARCO in clearing arthritogenic alphavirus particles from the circulation (Carpentier *et al*., 2019), we evaluated the extent to which the magnitude and duration of viremia is altered in MARCO^-/-^ mice. At 1-day post-inoculation (dpi), infectious virus in the blood of MARCO^-/-^ mice was elevated (230-fold; *P* < 0.0001) compared with WT mice (**Fig 1C**). Viremia peaked in both WT and MARCO^-/-^ mice at 2 dpi, but peak viremia was 150-fold higher in MARCO^-/-^ mice. In addition, MARCO^-/-^ mice maintained an elevated level of infectious virus in the serum through day five post-infection (**Fig 1C**). As neutralizing antibody can mask viral particles in the circulation at later times post-infection, we also analyzed serum samples for viral genomes by RT-qPCR. This analysis revealed that while CHIKV particles were mostly cleared from the blood of WT mice by 7 dpi, they remained detectable in the blood of MARCO^-/-^ mice (**Fig 1D**). These findings demonstrate that in the absence of MARCO both the magnitude and duration of viremia are increased.

We next evaluated whether the presence or absence of MARCO influences viral dissemination by quantifying viral burden in tissues proximal and distal to the inoculation site at 1 dpi. CHIKV burdens were similar in the ipsilateral ankle of WT and MARCO^-/-^ mice, suggesting that MARCO does not influence replication near the site of inoculation (**Fig 1E**). In contrast, MARCO^-/-^ mice had 2-3 orders of magnitude more infectious virus in distal tissues compared with WT mice (contralateral ankle: 3,163-fold, *P* < 0.0001; contralateral quadriceps: 951-fold, *P* < 0.0001) (**Fig 1E**). These findings were not unique to CHIKV, as infection of MARCO^-/-^ mice with two other arthritogenic alphaviruses, ONNV and RRV, also resulted in elevated viral burdens in distal tissues and in the serum at 1 dpi compared with WT mice (**Fig 1F-G**). The elevated viral burden observed in the contralateral ankle and quadriceps of MARCO^-/-^ mice infected with CHIKV persisted throughout the course of infection, with increased viral burden observed at days 3, 7 and 14 post-infection (**Fig S1A-C**). These findings demonstrate that in the absence of MARCO, arthritogenic alphaviruses develop a much higher viremia and are better able to disseminate to distal tissues.

### Kupffer cells rapidly remove CHIKV particles from the circulation

In prior studies, we found that i.v. inoculated CHIKV particles are rapidly cleared from the circulation and accumulate in the liver in a MARCO-dependent manner (Carpentier *et al*., 2019). Moreover, depletion of phagocytic cells in the spleen and liver in contact with the blood, through i.v. administration of clodronate-loaded liposomes (CLL), prevented accumulation of CHIKV RNA in the liver (Carpentier *et al*., 2019). These data suggest that KCs play a dominant role in the removal of CHIKV particles from the circulation. Based on these data, we hypothesized that MARCO-mediated clearance of viral particles by KCs limits the magnitude and duration of viremia, thus restricting viral dissemination and pathogenicity. To test this, we evaluated the extent to which specific depletion of KCs impaired the clearance of circulating CHIKV particles. To do this, we used Clec4F-DTR mice (Scott *et al*, 2016), which express the diphtheria toxin receptor (DTR) under the control of *Clec4f*, a gene expressed exclusively in KCs. We treated WT or Clec4F-DTR^+^ mice with DT 1-2 days prior to i.v. inoculation of CHIKV. In WT mice, CHIKV particles were rapidly cleared from the circulation by 45 min (**Fig 2A**). In contrast, Clec4F-DTR^+^ mice had increased viral particles in the serum at this time point (77-fold; *P* < 0.0001) (**Fig 2A**), demonstrating that KCs contribute to CHIKV clearance. However, the block to clearance in DT-treated Clec4F-DTR^+^ mice was not as robust as observed in mice treated i.v. with CLL (**Fig 2B**). Given the broad effects of CLL treatment (Seiler *et al*, 1997; Van Rooijen & Sanders, 1994), we employed CD169-DTR^+^ mice (Miyake *et al*, 2007), which allow for specific depletion of CD169^+^ cells, including liver KCs and splenic MMM and MZM (Gupta *et al*, 2016; Miyake *et al*., 2007). CHIKV clearance was blocked in DT-treated CD169-DTR^+^ mice to levels similar to that observed in CLL-treated mice (**Fig 2C**). However, we found that KC depletion was much more efficient in CLL-treated WT mice and DT-treated CD169-DTR^+^ mice compared with DT-treated Clec4F-DTR^+^ mice, with averages of 1.25 (CLL), 2.5 (CD169-DTR^+^) and 11.5 (Clec4F-DTR^+^) F4/80^+^ cells per field of view (**Fig 2D-E)**. The less efficient depletion of KCs in Clec4F-DTR^+^ mice likely accounts for the less potent block to clearance of circulating CHIKV particles. Collectively, these findings suggest that KCs are primarily responsible for the rapid removal of CHIKV from the circulation following i.v. inoculation.

**Figure 2.**
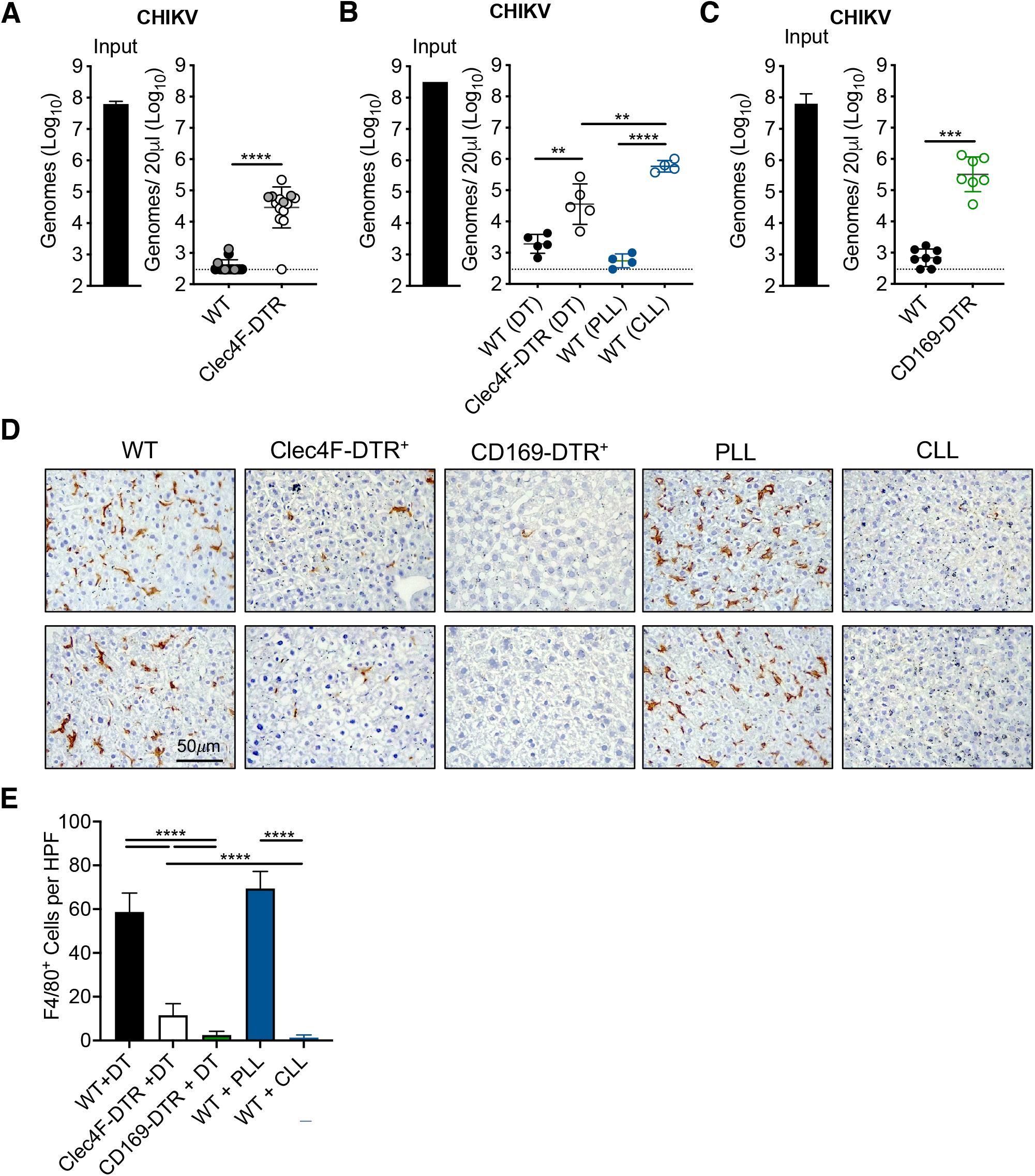
Depletion of Kupffer cells impedes CHIKV clearance from the circulation. (**A)** WT and Clec4F-DTR^+^ C57BL/6 mice were treated with diphtheria toxin (DT) either intraperitoneally (i.p.) 24 h prior to inoculation (n=10, two experiments, black dots) or intravenously (i.v.) 48 and 24 h prior to inoculation (n=4, one experiment, grey dots) to deplete KCs and inoculated i.v. with 10^8^ CHIKV particles. Viral genomes in the inoculum and serum at 45 minutes (min)-post inoculation were determined by RT-qPCR. Mean ± SD. Three experiments, n= 14. Mann-Whitney test; *\*\*\*\*P <* 0.0001. (**B**) WT or Clec-4F-DTR^+^ C57BL/6 mice were treated i.v. with DT, PLL or CLL prior to i.v. inoculation of 10^8^ CHIKV particles. Viral genomes were quantified as in (A). Mean ± SD. One experiment, n=4. Mann-Whitney test; \*\**P* < 0.01, \*\*\*\**P* < 0.0001. (**C**) WT or CD169-DTR^+^ mice were treated i.p. with DT prior to i.v. inoculation of 10^8^ CHIKV particles. Viral genomes were quantified as in (A). Mean ± SD. Two experiments, n=7-8. Mann-Whitney test; \*\*\**P <* 0.001. (**D**) Livers from WT, Clec4F-DTR^+^ or CD169-DTR^+^ mice treated i.p. with DT, or WT mice treated i.v. with PLL or CLL were analyzed by IHC to visualize F4/80^+^ macrophages (brown). Representative sections are shown. (**E**) F4/80^+^ cells in 10 randomly selected high-powered fields (HPF) per liver section were counted to calculate average F4/80^+^ cells per HPF. Mean ± SD. N=2-4 mice per group.

### Depletion of KCs is not sufficient to enhance early CHIKV dissemination

We reasoned that if clearance of viral particles by MARCO-expressing KCs was responsible for the enhanced viremia and dissemination observed in MARCO^-/-^ mice following s.c. virus inoculation, depletion of KCs would also enhance CHIKV dissemination. To test this idea, we treated WT or Clec4F-DTR^+^ mice with DT prior to s.c. inoculation of CHIKV in the left rear footpad and measured viral burden in tissues at 1 dpi. Surprisingly, unlike what we observed in MARCO^-/-^ mice (**Fig 1E**), the amounts of CHIKV in tissues and the circulation of WT and DT-treated Clec4F-DTR^+^ mice were indistinguishable (**Fig 3A**). Given that depletion of KCs is not complete in DT-treated Clec4F-DTR^+^ mice (**Fig 2D-E**), we i.v. treated WT mice with PLL or CLL 42 h prior to s.c. inoculation of CHIKV in the left rear footpad, as i.v. CLL treatment efficiently depletes F4/80^+^ cells in the liver (**Fig 2D-E**). Despite this, we found that CHIKV levels in distal tissues at 1 dpi were indistinguishable among PLL- and CLL-treated mice (**Fig 3B**). However, viremia was elevated in CLL-treated mice (27-fold; *P* < 0.001) (**Fig 3B**), confirming a role for phagocytic cells in limiting CHIKV viremia. These data demonstrate that while the absence of MARCO is sufficient to enhance CHIKV dissemination, depletion of KCs is not, suggesting that additional MARCO expressing cells restrict CHIKV dissemination.

**Figure 3.**
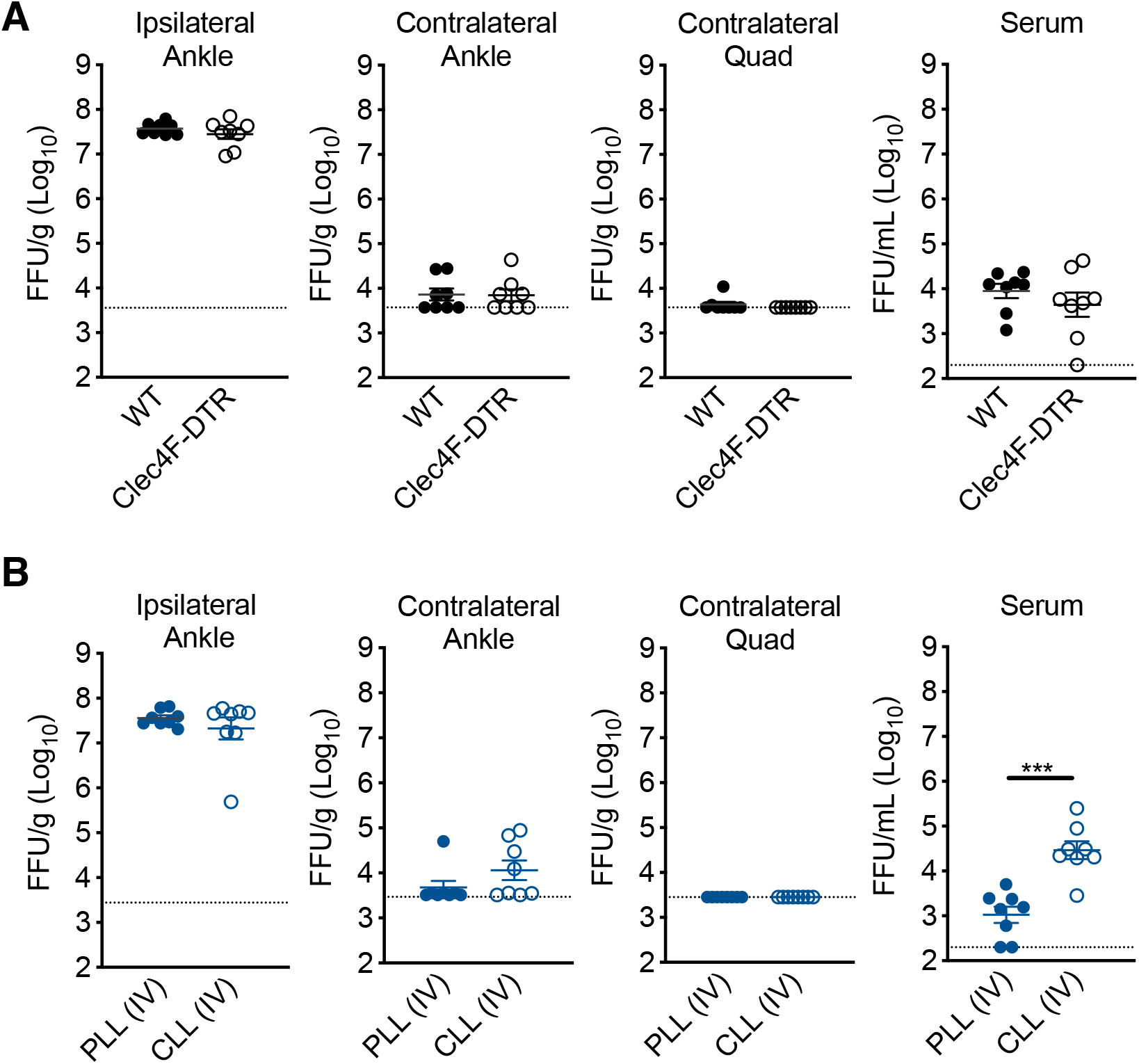
Depletion of KCs is not sufficient to enhance early CHIKV dissemination. (**A**) WT or Clec4F-DTR^+^ C57BL/6 mice were treated i.v. with DT prior to s.c. inoculation in the left rear footpad with 10^3^ PFU of CHIKV. Infectious virus was quantified at 24 hpi by FFA. Mean ± SEM. Two experiments, n=8. Mann-Whitney test; *P* > 0.05. (**B**) WT C57BL/6 mice were treated i.v. with PLL or CLL 42 h prior to s.c. inoculation in the left rear footpad with 10^3^ PFU of CHIKV. Infectious virus at 24 hpi was quantified by FFA. Mean ± SD. Two experiments, n=8. Mann-Whitney test; *\*\*\*P <* 0.001.

### The draining lymph node limits arthritogenic alphavirus dissemination

Following replication at the site of inoculation, arthritogenic alphaviruses spread through the lymph to the dLN before establishing viremia and disseminating to distal tissues. Given this, we hypothesized that MARCO-expressing cells in the dLN capture CHIKV particles, delaying the establishment of viremia and limiting viral dissemination to distal tissues. To test this, we used lymphotoxin alpha deficient mice (LT*α*^-/-^), which developmentally lack peripheral lymph nodes (De Togni *et al*, 1994). To evaluate the relative contributions of the dLN and liver in controlling CHIKV dissemination, WT or LT*α*^-/-^ mice were treated i.v. with PLL or CLL to deplete phagocytic cells in the liver prior to s.c. inoculation of CHIKV in the left rear footpad. At 1 dpi, the viral burden in tissues proximal and distal to the site of inoculation were quantified. In WT mice, low levels of virus were detected in distal tissues and serum of both PLL- and CLL-treated mice (**Fig 4A**). In contrast, PLL-treated LT*α*^-/-^ mice, which lack LNs but retain liver KCs, had an elevated viral burden in distal muscle (29-fold; *P* < 0.0001) and joint tissue (107-fold; *P* < 0.0001), and had moderately elevated viremia (10-fold; P < 0.0001) (**Fig 4A**). Compared with PLL-treated LT*α*^-/-^ mice, LT*α*^-/-^ mice treated with CLL, which lack both LNs and KCs, had higher viral burdens in distal muscle (4-fold; *P* < 0.05) and joint tissue (7-fold; *P* < 0.0001), and a highly elevated viremia (223-fold; *P* < 0.0001) (**Fig 4A**). These findings demonstrate that the dLN functions as a major barrier to CHIKV dissemination, as in its absence viral dissemination to distal tissues is strongly increased. The modest elevation in viremia in PLL-treated LT*α*^-/-^ mice compared with CLL-treated LT*α*^-/-^ mice suggests that phagocytic cells in the liver remove much of the circulating virus. An important role for liver phagocytes in controlling dissemination is supported by the elevated viral burden in both distal tissues and the circulation in CLL-treated LT*α*^-/-^ mice, which lack both LNs and KCs (**Fig 4A**).

**Figure 4.**
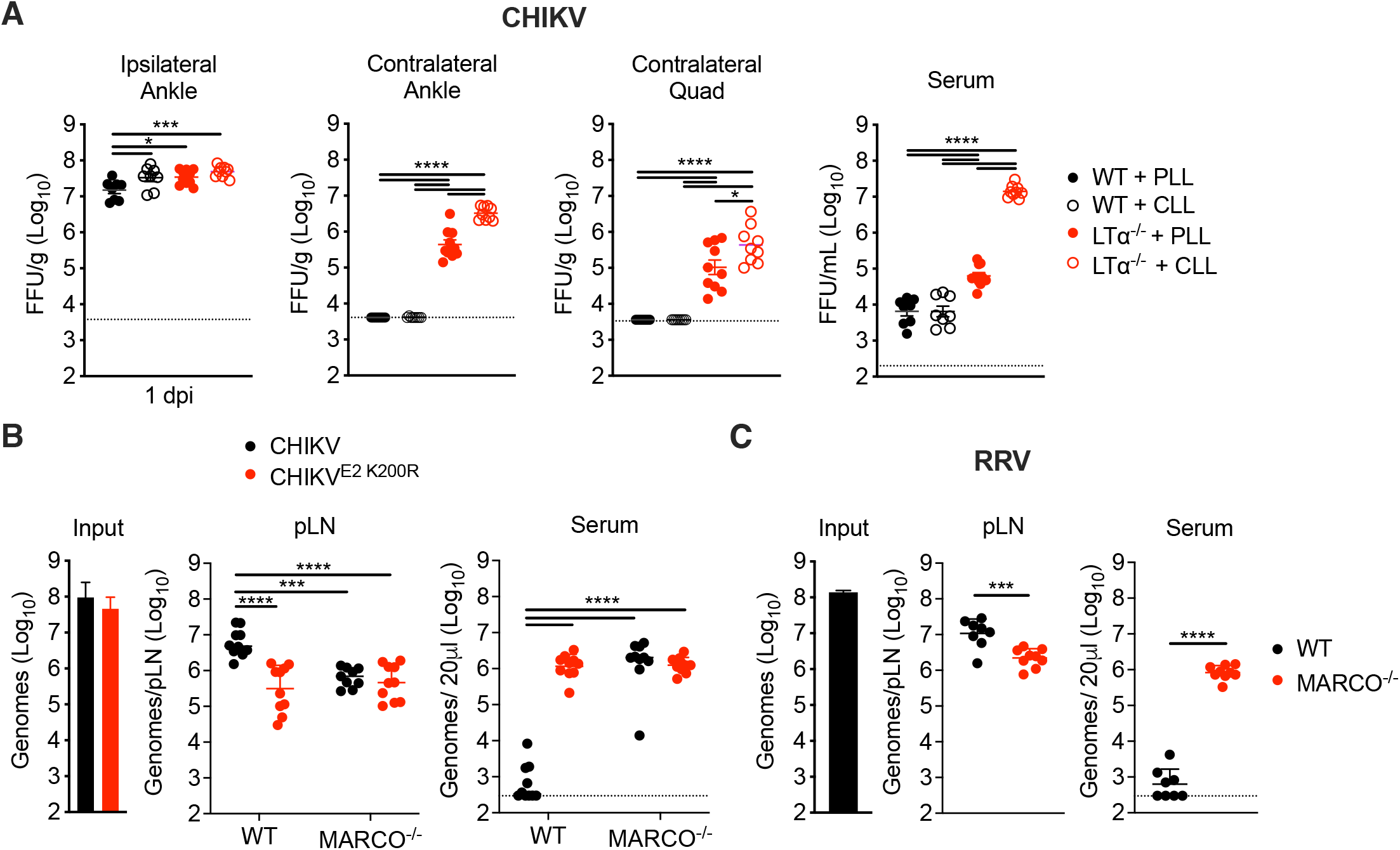
**The draining lymph node limits arthritogenic alphavirus dissemination. (**A) WT or LT*α*^-/-^ mice were treated i.v. with PLL or CLL 42 h prior to s.c. inoculation of the left rear footpad with 10^3^ PFU of CHIKV. Infectious virus at 24 hpi was analyzed by FFA. Mean ± SEM. Two experiments, n=8-10. Mann-Whitney test; \**P <* 0.05, *\*\*\*P <* 0.001, *\*\*\*\*P <* 0.0001. (**B**) WT or MARCO^-/-^ C57BL/6 mice were inoculated s.c. in the left rear footpad with 10^8^ particles of WT CHIKV or CHIKV E2 K200R. Viral genomes in the dLN and serum at 2 hpi were quantified by RT-qPCR. Mean ± SD. Two experiments, n=10. Two-way ANOVA with Bonferroni’s multiple comparisons test; \*\*\**P* < 0.001, \*\*\*\**P* < 0.0001. (**C**) WT or MARCO^-/-^ C57BL/6 mice were inoculated s.c. in the left rear footpad with 10^8^ particles of RRV. Viral genomes in the dLN and serum were quantified by RT-qPCR. Mean ± SD. Two experiments, n=8-9. Mann-Whitney test; \*\*\**P* < 0.001, \*\*\*\**P* < 0.0001.

The enhanced viral dissemination observed in CLL-treated LT*α*^-/-^ mice (**Fig 4A**) is similar to what was observed in MARCO^-/-^ mice (**Fig 1E**), suggesting that MARCO-expressing cells in the dLN sequester alphavirus particles. To explore this idea, we inoculated WT or MARCO^-/-^ mice s.c. in the footpad with WT CHIKV or CHIKV E2 K200R (mutant virus that evades MARCO). At 2 hpi, a time point at which no new infectious virus has been produced, we collected the dLN and serum to evaluate the fate of the inoculated virus. In WT mice inoculated with WT CHIKV, we detected high levels of viral RNA in the dLN, while little to no virus was observed in the serum (**Fig 4B**). In contrast, WT mice inoculated with CHIKV E2 K200R and MARCO^-/-^ mice inoculated with either WT CHIKV or CHIKV E2 K200R had 7-10-fold lower levels of viral RNA in the dLN, and remarkably had 3,700-6,200-fold more virus in the serum (**Fig 4B**). Moreover, we found that MARCO^-/-^ mice inoculated with RRV, a closely related arthritogenic alphavirus, also had reduced viral RNA in the dLN (5-fold; *P* < 0.001) and higher levels of virus in the blood (1,790-fold; *P* < 0.0001) compared with WT mice (**Fig 4C**). Importantly, MARCO^-/-^ mice inefficiently clear circulating viral particles (Carpentier *et al*., 2019). Therefore, the increased viremia in MARCO^-/-^ mice 2 h following s.c. virus inoculation is likely a reflection of not only reduced virus accumulation in the dLN, but also the lack of clearance of circulating virus by liver KCs.

### LN Macrophages are not required for CHIKV accumulation in the dLN or for limiting viral dissemination

We next sought to define the cell type(s) in the dLN that limit CHIKV dissemination. Within the LN, MARCO is reported to be expressed by MS macrophages (Elomaa *et al*, 1995) and a subset of LECs in the medullary region (Fujimoto *et al*, 2020; Takeda *et al*, 2019; Walsh *et al*, 2021; Xiang *et al*, 2020). Consistent with these data, using flow cytometry we found that MARCO was expressed specifically on MS macrophages and LECs (**Fig S2**). Previous studies found that macrophages in the dLN capture lymph-borne viruses (Farrell *et al*., 2015; Gonzalez *et al*, 2010; Hickman *et al*, 2008; Junt *et al*., 2007), and that CLL-mediated depletion of dLN macrophages decreased viral capture by the dLN and increased viremia and dissemination (Farrell *et al*., 2015; Junt *et al*., 2007). Given this, we hypothesized that MARCO^+^ MS macrophages limit the dissemination of arthritogenic alphaviruses. As MS and SCS macrophages in the LN are CD169^+^ (Gray & Cyster, 2012), we used CD169-DTR mice to deplete LN macrophages and evaluate their capacity to promote virus accumulation in the dLN and limit virus accumulation in the circulation. WT or CD169-DTR mice were treated with DT, which results in efficient depletion of CD169^+^ cells in the dLN (**Fig 5A**). Mice were then inoculated with CHIKV s.c. in the footpad and at 2 hpi, viral RNA in the dLN and serum was quantified by RT-qPCR. Remarkably, there was no difference in the amount of viral RNA detected in the dLN of DT-treated WT or CD169-DTR^+^ mice, and low levels of viral RNA in the blood were observed in both WT and CD169-DTR^+^ mice (**Fig 5B**). Importantly, DT treatment of CD169-DTR mice also depletes KCs (**Fig 2D** and **3E**), and thus any virus that traffics from the dLN to the blood should remain circulating and not be masked by the clearance effects of KCs. These findings suggest that macrophages in the dLN are not required to sequester viral particles and limit access to the circulation. Consistent with this, CHIKV dissemination to distal tissues was not enhanced in DT-treated CD169-DTR^+^ mice compared with WT mice (**Fig 5C**), or in mice treated with CLL both i.v. and s.c. in the footpad to deplete phagocytic cells in the liver and dLN, respectively (**Fig S3**). These findings reveal that in contrast to previous reports for other viruses (Junt *et al*., 2007), LN macrophages are not required for clearance of arthritogenic alphavirus particles from the lymph.

**Figure 5.**
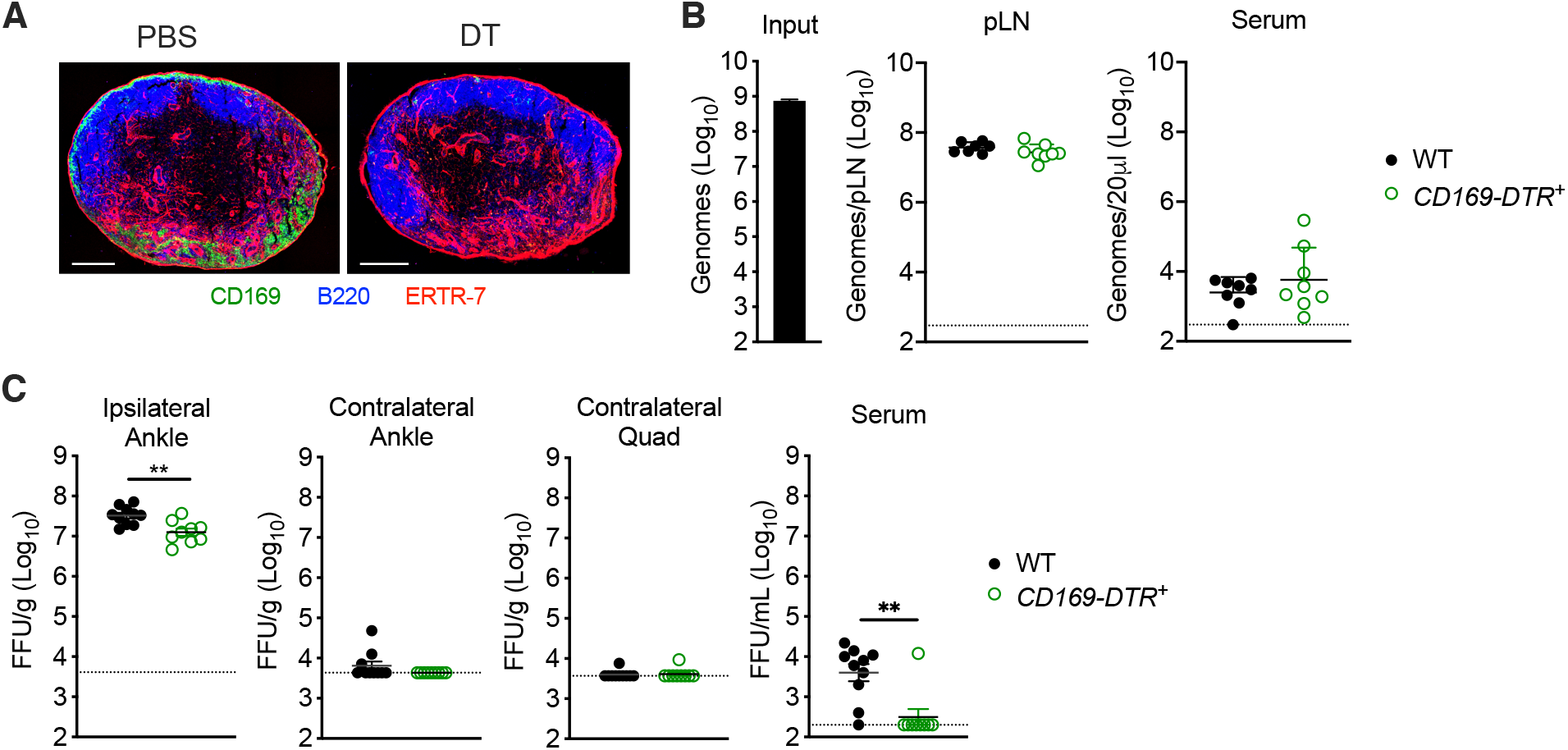
Macrophages in the dLN are not required for CHIKV accumulation in the dLN or limiting viral dissemination. (**A**) CD169-DTR^+^ mice were treated i.p. with PBS or DT prior to collection of the popliteal LN. Frozen LN sections were stained for CD169^+^ macrophages (green), B220^+^ B cells (blue) or ERTR-7^+^ stromal cells (red). (**B**) WT or CD169-DTR^+^ C57BL/6 mice were treated i.p. with DT prior to s.c. inoculation in the left rear footpad with 10^8^ particles of WT CHIKV. Viral genomes in the dLN and serum at 2 hpi were quantified by RT-qPCR. Mean ± SD. Two experiments, n=8. Mann-Whitney test; *P* > 0.05. (**C**) WT or CD169-DTR^+^ C57BL/6 mice were treated i.p. with DT prior to s.c. inoculation in the left rear footpad with 10^3^ PFU of CHIKV. Infectious virus at 24 hpi was quantified by FFA. Mean ± SEM. Two experiments, n=9-10. Mann-Whitney test; \*\**P* < 0.01.

### CHIKV-E2-mCherry particles co-localize with MARCO^+^ LECs in the dLN

Given that medullary macrophages are not required for accumulation of arthritogenic alphavirus particles in the dLN, we next investigated the role of LECs, as recent reports identified a subset of LECs in the medullary region of the lymph node that express MARCO (Fujimoto *et al*., 2020; Xiang *et al*., 2020). Moreover, LECs have been shown to capture and archive viral antigen (Tamburini *et al*, 2014), further supporting a potential role for LECs in capturing alphavirus particles. To determine whether viral particles colocalized with MARCO^+^ LECs in the dLN, we used a recombinant CHIKV in which mCherry is fused to the E2 glycoprotein present in viral particles (CHIKV-E2-mCherry). Importantly, CHIKV-E2-mCherry particles were efficiently cleared from the circulation, and clearance could be blocked by pre-treatment of mice with poly(I), a class A SR inhibitor that competitively inhibits MARCO (Chen *et al*, 2006) (**Fig S4**). CHIKV-E2-mCherry particles were inoculated s.c. into the foot and the popliteal dLN was collected at 2 hpi. Frozen LN sections were stained for Lyve1^+^ LECs, MARCO, B220 (marking B cell follicles and revealing nodal orientation) and mCherry^+^ CHIKV particles. As previously reported, we detected a robust MARCO^+^ Lyve1^+^ LEC population in both infected and uninfected WT mice that was primarily restricted to the LN medullary sinus (**Fig 6A**). Although MARCO^-/-^ mice lacked MARCO^+^ cells as expected, the medullary sinus remained intact with similar morphology to that of WT mice (**Fig 6A**, middle panels). Visually, mCherry staining for CHIKV particles was most intense in Lyve1^+^ MARCO^+^ LECs (**Fig 6A**, far right panels). To better quantify CHIKV^+^ cells in the LECs of WT and MARCO^-/-^ mice, we first colocalized mCherry and Lyve1^+^ signals in the LN, detecting many double positive cells in WT but not MARCO^-/-^ LNs (**Fig 6B-C**). Indeed, while 39.3% of Lyve1^+^ voxels in were also mCherry^+^ in WT LNs, only 3.0 % were mCherry^+^ in MARCO^-/-^ mice, consistent with the decreased accumulation of viral genomes in the dLN of MARCO^-/-^ mice at 2 hpi (**Fig 5B**). Higher magnification images demonstrated that in WT mice, mCherry staining overlapped with both Lyve1 and MARCO staining (**Fig 6D**). In contrast, in MARCO^-/-^ LNs, we observed only a small amount of mCherry staining in CD11b^+^ cells, some of which co-expressed CD169 (representing medullary sinus macrophages) (**Fig 6E**). Collectively, these findings demonstrate that following s.c. inoculation, CHIKV particles are bound by medullary sinus LECs using MARCO^-/-^.

**Figure 6.**
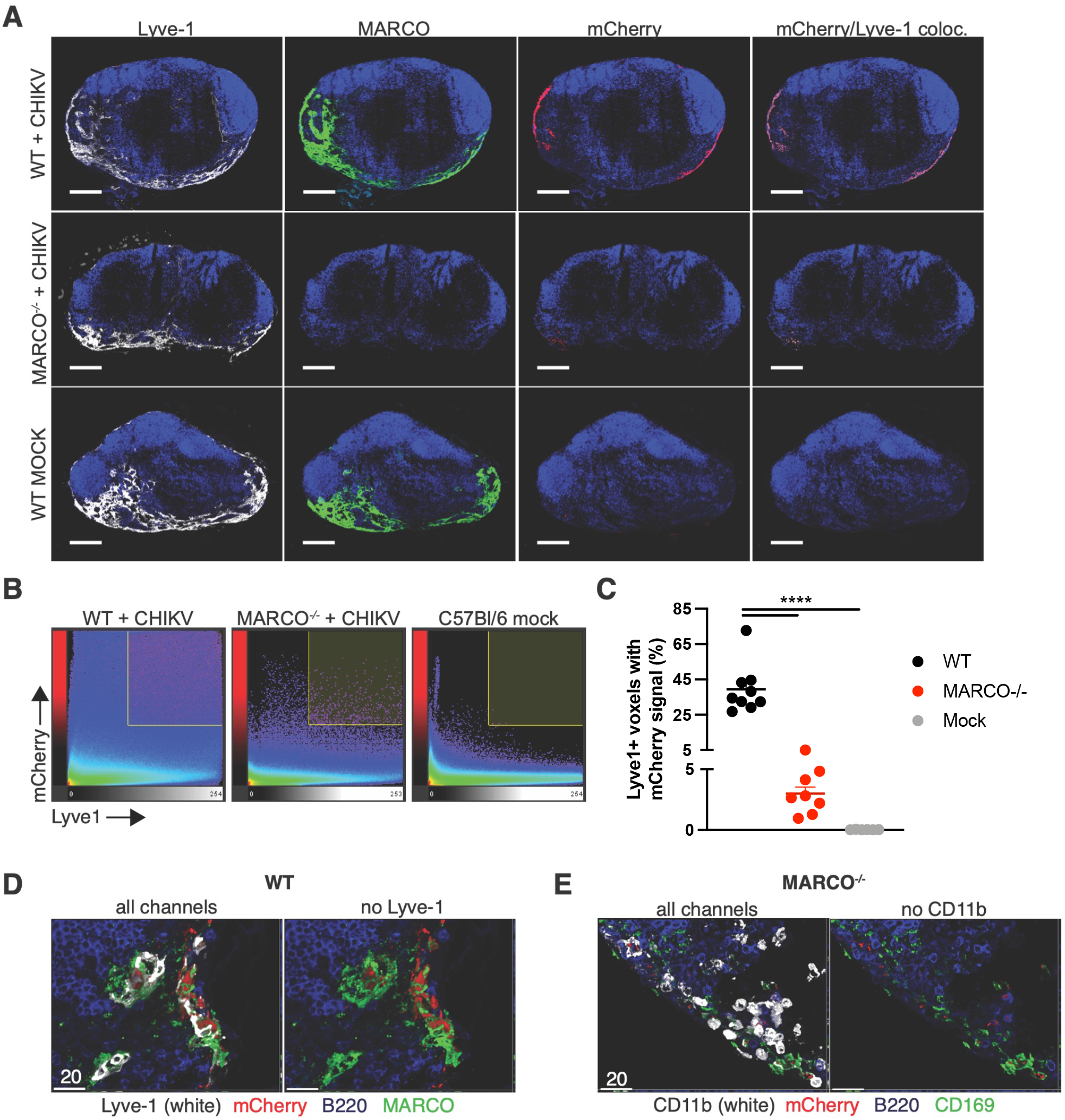
MARCO^+^ LECs capture lymph-borne CHIKV particles. WT or MARCO^-/-^ C57BL/6 mice were inoculated s.c. in the rear feet with 5*10^4^ PFU of CHIKV-E2-mCherry and the popliteal dLNs were collected at 2 hpi. **(A)** Frozen dLN sections were stained for Lyve-1^+^ LECs (white), MARCO (green), B220 (blue) and mCherry CHIKV particles (red). **(B)** Representative histograms of confocal images showing voxel intensities for Lyve1 and mCherry. **(C)** Percent of Lyve1^+^ voxels with mCherry signal; dots represent individual sections. **(D)** Higher magnification confocal image of a WT lymph node section stained as in (A) (**E**) Higher magnification confocal image of a MARCO^-/-^ lymph node stained for CD11b^+^ cells (white), mCherry^+^ CHIKV particles (red), B220 B cells (blue), and CD169 macrophages (green). Data are representative of 2 experiments, n=6-9. One-way ANOVA with Tukey’s multiple comparisons test; \*\*\*\**P*<0.0001.

### MARCO^+^ LECs harbor CHIKV RNA

To further characterize LEC subsets that capture virus, at 24 hpi we generated single cell suspensions from the dLN for three biological replicates each of mock- and CHIKV-infected mice. We then enriched for CD45^-^ cells (**Fig S5**) and performed scRNA-seq to identify cell populations that harbor viral RNA. Between mock- and CHIKV-infected samples, which clustered distinctly, we captured a total of 60,185 cells (**Fig 7A**). To identify the cell types represented in these samples, we used an automated approach (Fu *et al*, 2020) that classifies cells based on their correlation with reference RNA-seq data. Using published data for cell types found in the mouse LN (Heng, 2008; Malhotra *et al*, 2012; Rodda *et al*, 2018), we were able to identify large populations of endothelial cells, non-endothelial stromal cells including fibroblastic reticular cells (FRC) and perivascular cells (PvC), along with smaller populations of CD45^+^ cells including B cells, T cells, and macrophages (**Fig 7B**). To identify endothelial cell subsets, we further divided the endothelial cells into niche-specific subpopulations using published reference data (Xiang *et al*., 2020). By this method, we identified blood endothelial cells (BEC) as well as LEC subsets including MARCO^+^, ceiling (cLEC), floor (fLEC), valve, collecting, Ptx3, and transition zone (tzLEC) (**Fig 7C and EV6**). Cells collected from the dLN of CHIKV-infected mice contained fewer LEC subsets and were mainly composed of MARCO^+^ LECs, valve LECs, and a population of endothelial cells that we were unable to further classify (**Fig 7C**). The limited number of LEC subsets identified in CHIKV-infected samples could be due to cell death within the dLN at 24 hpi. However, one caveat to our approach is that we are using reference data from uninfected mice which could make it more challenging to accurately annotate LN cell populations during CHIKV infection.

**Figure 7.**
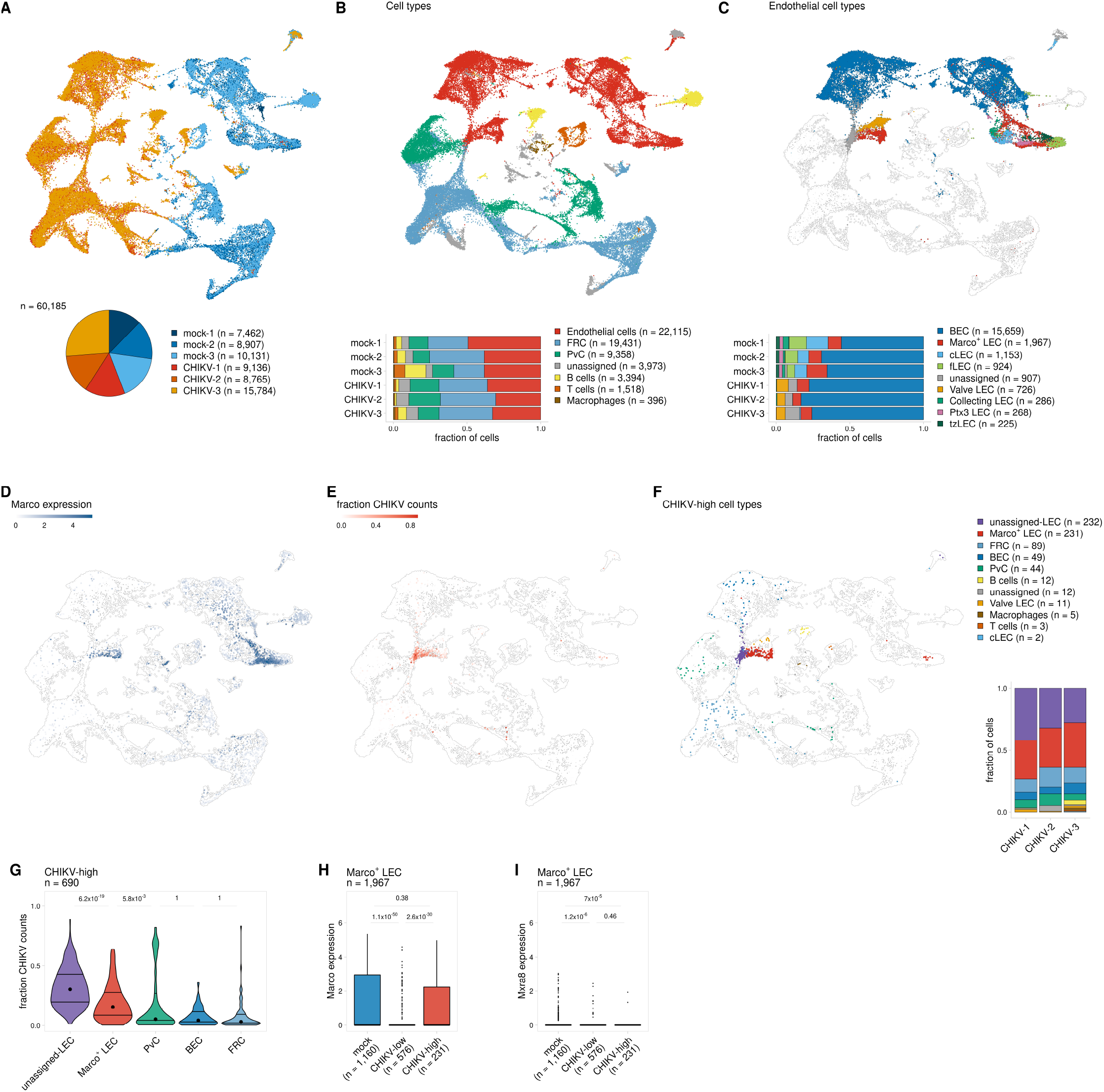
MARCO^+^ LECs harbor CHIKV RNA. WT C57BL/6 mice were s.c. inoculated with PBS (mock, n=3) or 10^3^ PFU of CHIKV (n = 3) in the left rear footpad. At 24 hpi, the dLN was collected and enzymatically digested into a single cell suspension. Cells were enriched for CD45- cells and analyzed by scRNA-seq as described in the materials and methods. (**A**) UMAP projection shows each replicate for mock- and CHIKV-infected mice; the number of cells obtained for each replicate is shown at the bottom. (**B**) UMAP projection shows annotated cell types (top) and the proportion of cells identified for each cell type (bottom). (**C**) UMAP projection shows endothelial subtypes (top) and the proportion of cells identified for each cell type (bottom). Non-endothelial cells are shown in white. (**D**) UMAP projection shows Marco expression. (**E**) UMAP projection shows the fraction of counts that align to the CHIKV genome. (**F**) UMAP projection shows cell types for cells classified as CHIKV-high. CHIKV-low cells are shown in white. The proportion of CHIKV-high cells belonging to each cell type is shown on the right. (**G**) The fraction of counts that align to the CHIKV genome is shown for CHIKV-high cells. Only cell types that include >20 cells are shown. (**H, I**) MARCO (**H**) and Mxra8 (**I**) expression is shown for MARCO^+^ LECs for mock-infected cells and CHIKV-infected cells classified as either CHIKV-low or CHIKV-high. *P*-values were calculated using a two-sided Wilcoxon rank-sum test with Bonferroni correction.

To identify cell types harboring CHIKV RNA, we classified cells based on the number of viral sequence counts. As expected, we detected only background levels of CHIKV RNA (3/26,500 cells with 1 CHIKV count each) in dLN cells of mock-infected mice (**Fig S7A**), while cells collected from the dLN of CHIKV-infected mice had viral RNA sequence counts as high as >8,000 per cell (**Fig S7A**). We used k-means clustering to divide cells from each sample into CHIKV-low and CHIKV-high groups and identified a small number of cells (n = 690, 1.1%) with high amounts of CHIKV RNA (**Fig S7A**). CHIKV-high cells displayed fewer mouse mRNA counts per cell and fewer expressed mouse genes (**Fig S7B**) potentially due to inhibition of host cell transcription or cell lysis, both of which can occur in CHIKV-infected cells (Fros & Pijlman, 2016).

We next analyzed the cell populations containing high amounts of CHIKV RNA. CHIKV-high cells are mainly composed of MARCO^+^ LECs (n = 231, 33%), FRCs (n = 89, 13%), BECs (n = 49, 7%), and PvCs (n = 44, 6%), along with a group of LECs that we were unable to further classify (unassigned-LEC, n = 232, 34%) (**Fig 7F**). Among the CHIKV-high cell types, we found that the unassigned-LECs and MARCO^+^ LECs show the highest viral burden as indicated by a high fraction of CHIKV counts per cell (**Fig 7E-G**), suggesting that these are the predominant cell populations in the dLN that capture viral particles. To further investigate the role of MARCO in these interactions, we compared MARCO expression for CHIKV-low and CHIKV-high MARCO^+^ LECs. This analysis revealed that CHIKV-high MARCO^+^ LECs had significantly higher expression of MARCO in comparison to their CHIKV-low counterparts (**Fig 7D** and **7H**). Moreover, CHIKV-low and CHIKV-high MARCO^+^ LECs expressed little to no Mxra8, a cell entry receptor for CHIKV (Zhang *et al*, 2018) (**Fig 7I**). These data further support a role for MARCO and LECs in sequestering viral particles in the dLN and limiting CHIKV dissemination.

## Discussion

Our results reveal a critical role for the scavenger receptor MARCO in controlling arthritogenic alphavirus viremia, dissemination, and disease. Similar protective roles for MARCO have been observed during other infections. For example, MARCO^-/-^ mice are impaired in their ability to clear *Streptococcus pneumonia* from the nasopharynx and lungs (Arredouani *et al*, 2004; Dorrington *et al*, 2013). Moreover, MARCO enhances phagocytosis of *Mycobacterium tuberculosis in vitro* (Bowdish *et al*, 2009), and MARCO polymorphisms are associated with altered susceptibility to pulmonary tuberculosis (Bowdish *et al*, 2013; Lao *et al*, 2017; Ma *et al*, 2011; Thuong *et al*, 2016). Finally, during influenza A virus infection in mice, MARCO suppresses early immunopathologic inflammatory responses, and accordingly, MARCO^-/-^ mice have increased morbidity and mortality compared with WT mice (Ghosh *et al*, 2011). However, MARCO also can be exploited by pathogens. For example, herpes simplex virus 1 (HSV-1) interactions with MARCO enhance epithelial cell adsorption, and MARCO^-/-^ mice have reduced wound sizes following s.c. HSV-1 inoculation (MacLeod *et al*, 2013).

Our findings identify two distinct MARCO expressing cell types that limit arthritogenic alphavirus dissemination and viremia: MARCO^+^ LECs in the dLN and KCs in the liver. KCs are well established to play a critical role in controlling bacteremia (Jenne & Kubes, 2013; Lee *et al*., 2010) However, the role of KCs in controlling viremia is not as well characterized. We find that specific depletion of KCs using Clec4F-DTR^+^ mice impairs CHIKV clearance from the circulation. While it remains possible that MARCO^+^ MZM in the spleen contribute, our findings demonstrate that KCs are the major cell type involved in the efficient removal of arthritogenic alphavirus particles from the blood, expanding their surveillance function to arboviruses.

Despite the critical role for KCs in removing alphavirus particles from the circulation, we found that depletion of KCs had no impact on CHIKV dissemination following s.c. viral inoculation. This led us to investigate the role of MARCO^+^ cells in the dLN in controlling CHIKV viremia and dissemination, as lymph nodes can function as barriers to pathogen dissemination (Bogoslowski & Kubes, 2018). For example, within minutes of subcutaneous inoculation, fluorescently labeled vesicular stomatitis virus (VSV) particles can be found trapped within SCS macrophages in the dLN (Junt *et al*., 2007). This observation extends to other viruses, including adenovirus (AdV) and vaccinia virus (VV) (Hickman *et al*., 2008; Junt *et al*., 2007), thus lending to the description of these SCS macrophages as “molecular flypaper” in regard to their ability to capture incoming viral particles. This macrophage-mediated capture has important implications for pathogen dissemination, as depletion of macrophages in the draining lymph node via s.c. CLL administration decreased accumulation of VSV in the dLN at early times post-infection, and increased viral dissemination to the blood (Junt *et al*., 2007). Similarly, depletion of macrophages in the dLN was shown to enhance the dissemination of murine cytomegalovirus (MCMV), West Nile virus (WNV), and *Pseudomonas aeruginosa (P. aeruginosa)*, and facilitate CNS invasion of neurotropic VSV (Farrell *et al*., 2015; Iannacone *et al*., 2010; Kastenmuller *et al*., 2012; Winkelmann *et al*, 2014).

Our results demonstrate the dLN is a major barrier to arthritogenic alphavirus dissemination, but unlike prior reports of macrophage-mediated capture our findings uncover a previously unrecognized role for LECs in scavenging arboviral particles to impair dissemination. We found that CHIKV-E2-mCherry particles colocalized with MARCO^+^ LECs in the dLN, and scRNA sequencing of dLN stromal cell populations identified MARCO^+^ LECs as the predominant cell type harboring CHIKV RNA. Notably, CHIKV RNA levels among MARCO^+^ LECs correlated with MARCO expression levels, with CHIKV-high cells showing higher expression of MARCO, suggesting MARCO may mediate internalization. MARCO^+^ LECs were negative for *Mxra8*, a known arthritogenic alphavirus entry receptor (Zhang *et al*., 2018). The genetic absence of *Mxra8* in mice reduced but did not eliminate viral replication and dissemination *in vivo* (Zhang *et al*, 2019), demonstrating that additional entry receptors exist. Notably, MARCO has been reported to facilitate entry of other viruses, including HSV-1, VV, and adenovirus into target cells (MacLeod *et al*, 2015; MacLeod *et al*., 2013; Maler *et al*, 2017; Stichling *et al*, 2018).

The role of LECs in capturing lymph-borne viral particles likely extends beyond arthritogenic alphaviruses. While macrophages were reported to play a major role in capture of other viruses, fluorescently labeled VSV, AdV, and VV particles co-localized with LECs in the medullary region of the dLN (Junt *et al*., 2007; Reynoso *et al*, 2019), which is where MARCO^+^ LECs reside. Future investigations are needed to understand whether MARCO is responsible for broadly mediating capture of diverse viruses by LECs, or whether other pattern recognition receptors (PRRs) are also involved. LECs express a wide range of PRRs, including toll-like receptors, Fc receptors, C-type lectin receptors, and additional scavenger receptors, suggesting they may have multiple mechanisms for scavenging diverse viral particles (Berendam *et al*, 2019; Jalkanen & Salmi, 2020).

Additional work is also needed to better understand the consequences of viral capture by LECs. Our findings suggest that capture of arthritogenic alphaviruses by MARCO^+^ LECs contributes to the control of viral dissemination. However, whether LECs become productively infected by arthritogenic alphaviruses remains under investigation. In prior studies, we were unable to detect fluorescent signal in the dLN following s.c. inoculation with a recombinant CHIKV expressing the fluorescent protein mKate (McCarthy *et al*, 2018). However, our scRNA-seq results reveal that only a small fraction of stromal cells in the dLN harbor CHIKV RNA, suggesting flow cytometry may not be sensitive enough to detect whether the virus is productively replicating in these cells. Our scRNA-seq analysis provides hints that the MARCO^+^ LECs may be actively infected. For example, cells harboring CHIKV RNA have high viral reads, suggestive of genome replication, and CHIKV-high cells have reduced reads for mouse genes, which is consistent with virus-mediated transcriptional shutoff (Fros & Pijlman, 2016). Further studies are needed, but these findings raise the possibility that MARCO facilitates arthritogenic alphavirus entry and infection of distinct cell types, such as MARCO^+^ LECs.

LEC-mediated capture of viral particles also could influence innate and adaptive immune responses. LECs have been reported to archive viral antigen for weeks following the resolution of the adaptive immune response (Kedl *et al*, 2017; Tamburini *et al*., 2014), and our findings may provide insight as to how these antigens are initially acquired by LECs. This archived antigen can be either directly presented or exchanged with dendritic cells to allow for cross-presentation to CD8^+^ T-cells to stimulate memory T cells and augment protective immunity (Kedl *et al*., 2017; Tamburini *et al*., 2014; Vokali *et al*, 2020). In other studies evaluating factors that influence alphavirus viremia and dissemination, injection of Semliki Forest virus, a closely related alphavirus, at the site of a mosquito bite in the skin of mice was found to delay viral spread to the lymph node, which ultimately enhanced early viremia and viral dissemination, and led to more severe disease outcomes (Pingen *et al*, 2016). It is possible that the retention of viral particles at the site of inoculation allows the virus to replicate to high titers before initiating potent immune responses due to viral capture in the dLN. Future studies are necessary to better understand how MARCO^+^ LEC-mediated capture of arthritogenic alphaviruses influences downstream innate and adaptive immune responses.

In summary, our results reveal a critical scavenging role for MARCO during arthritogenic alphavirus infection. We find that following s.c. inoculation, alphavirus particles accumulate in the dLN in association with MARCO^+^ LECs, limiting viral spread to the blood. Once reaching the blood, liver KCs provide a second line of defense and rapidly clear circulating alphavirus particles in a MARCO-dependent manner. Collectively, these findings advance our mechanistic understanding of how viremia is controlled during arboviral infections, which has several important implications for arboviral biology. First, viremia has been shown to positively correlate with disease severity following infection with CHIKV and other arboviruses (Chow *et al*., 2011; de St Maurice *et al*., 2018; Pozo-Aguilar *et al*., 2014; Vaughn *et al*., 2000; Vuong *et al*., 2020; Waggoner *et al*., 2016). Consistent with this, we find that the high magnitude and duration of viremia observed in CHIKV-infected MARCO^-/-^ mice promoted more rapid viral dissemination, increased viral tissue burdens, and resulted in more severe disease signs. These finding raise the possibility that MARCO also influences disease severity in humans. The MARCO gene is highly polymorphic in humans and mice (Bowdish & Gordon, 2009), and this genetic variation has been demonstrated to influence human susceptibility to tuberculosis and respiratory syncytial virus (Bowdish *et al*., 2013; High *et al*, 2016; Lao *et al*., 2017; Ma *et al*., 2011; Thuong *et al*., 2016). Given our findings, it is possible that MARCO polymorphisms similarly influence disease severity following arthritogenic alphavirus infection. In addition to influencing disease severity, MARCO-virus interactions likely also affect virus transmission efficiency and reservoir host competency in nature, as the magnitude and duration of viremia is an important factor dictating which vertebrate species can serve as reservoirs for arboviruses (Weaver, 2018). Thus, differences in MARCO alleles may influence which vertebrate hosts participate in arthritogenic alphavirus transmission cycles. While humans are dead-end hosts for most arboviruses, a select few including CHIKV, DENV, and ZIKV generate a sufficiently high level of viremia to facilitate human-mosquito-human transmission cycles (Weaver, 2018). These viruses pose a high risk for emergence and re-emergence, as evidenced by the now global distribution of DENV and the recent ZIKV and DENV epidemics. This underscores the need for an improved understanding of viremic control. Collectively, our findings shed light on the mechanistic control of viremia during arboviral infections, and more broadly advance our understanding of how the lymph node restricts virus dissemination.

## Materials and Methods

### Ethics Statement

This study was performed in strict accordance with the recommendations in the Guide for the Care and Use of Laboratory Animals of the National Institutes of Health. All of the animals were handled according to approved institutional animal care and use committee (IACUC) protocols (#00026) of the University of Colorado School of Medicine (Assurance Number A3269-01). Experimental animals were humanely euthanized at defined endpoints by exposure to isoflurane vapors followed by thoracotomy.

### Cells

Vero cells (ATCC CCL81) were cultured at 37°C in Dulbecco’s Modified Eagle medium (DMEM)-F-12 (Gibco) supplemented with 10% fetal bovine serum, 1x nonessential amino acids (Life Technologies), and 1X penicillin-streptomycin. BHK-21 cells (ATCC CCL10) were cultured at 37°C in α-minimum essential medium (Gibco) supplemented with 10% FBS, 10% tryptone phosphate broth, and penicillin-streptomycin.

### Viruses

The CHIKV strain used in these studies is AF15561, an Asian genotype strain isolated from a human patient in Thailand (GenBank accession no. EF452493). cDNA clones of AF15561 and AF15561 E2 K200R have been described previously (Hawman *et al*., 2017). The recombinant CHIKV AF15561 cDNA clone encoding mCherry-tagged E2 glycoprotein was derived from a 181/25 CHIKV E2 mCherry-tagged cCNA clone kindly provided by Richard Kuhn (Purdue University). Site-directed mutagenesis was first used to revert positions E2 12 and E2 84 from attenuated 181/25 to WT AF15561 using the following primers: CHIKV-181/25 E2 I12T FOR (5′-gtgagctaggtacggtcttgtggctttatagacattgaa-3′), CHIKV-181/25 E2 I12T REV (5′-ttcaatgtctataaagccacaagaccgtacctagctcac-3′), CHIKV 181/25 E2 R82G FOR (5′-gttcttacaaatagcccggccctctctgcgtc-3′) and CHIKV 181/25 R82G Rev (5′-gacgcagagagggccgggctatttgtaagaac-3′). A fragment containing part of capsid, mCherry, and part of E2 was then subcloned into an AF15561 cDNA clone using restriction sites XhoI and XmaI. To generate virus stocks, linearized cDNA clones were *in vitro* transcribed with SP6 RNA polymerase, and viral RNA was electroporated into BHK-21 cells as described previously (Ashbrook *et al*, 2014). At 24-28 h post-electroporation, clarified supernatant containing infectious virus was collected, aliquoted and stored at -80°C. The RRV strain used is SN11 (Liu *et al*, 2011), a clinical isolate (kindly provided by John Aaskov, Queensland University of Technology) that was passaged 1X on C6/36 cells before we propagated the stock used in these studies in BHK-21 cells. ONNV SG650 (Lanciotti, 1998), a strain isolated from human sera in Uganda in 1996, was derived from a cDNA clone ((Vanlandingham, 2006); provided by Stephen Higgs, Kansas State University) through electroporation into BHK-21 cells, and propagated on BHK-21 cells for one passage to increase titer. Infectious virus was titered by plaque assay on BHK-21 cells. To quantify viral genomes, viral stocks were treated with RNase1 at 37°C for 1 h. RNA was extracted and viral genomes were quantified by RT-qPCR.

### Mouse Experiments

WT C57BL/6 and congenic Lymphotoxin alpha^-/-^ (LT*α*^-/-^) mice (De Togni *et al*., 1994) were obtained from the Jackson Laboratory. Congenic CD169-DTR^+^ (Miyake *et al*., 2007) mice were provided by Jason Cyster (University of California San Francisco) and congenic MARCO^-/-^ mice (Arredouani *et al*., 2004) were provided by Dawn Bowdish (McMaster University). Clec4F-DTR^+^ C57BL/6 mice (Scott *et al*., 2016) were provided by Martin Guilliams (Ghent University). CD169-DTR^+^, MARCO^-/-^, LT*α*^-/-^ and Clec4F-DTR^+^ mice were housed and bred at the University of Colorado School of Medicine under specific pathogen-free conditions and were distributed randomly into groups containing approximately even division of sexes for experiments. WT male mice were purchased commercially and were age matched and distributed randomly across groups. Mice 4 weeks of age were used in all experiments. All mouse experiments were performed under animal biosafety level 2 or 3 conditions, as appropriate.

For experiments involving Clec4F-DTR^+^ mice, mice were treated with 50 ng of DT either i.v. or i.p. as indicated in the Figure legend, 48 h and 24 h prior to virus inoculation. For experiments involving CD169-DTR^+^ mice, mice were injected with 100 ng of DT i.p. 48 h and 24 h prior to virus inoculation. For experiments involving depletion of liver and splenic phagocytes, mice were inoculated i.v. with 100 μl per 10 g of body weight of PBS-(PLL) or clodronate-loaded liposomes (CLL) (clodronateliposomes.org) 42 h prior to virus inoculation. To deplete phagocytic cells in the draining lymph node, mice were inoculated s.c. in the left-rear footpad with 20 μl of PLL or CLL 24 h prior to virus inoculation.

In experiments evaluating disease or viral tissue burdens, mice were anesthetized with isoflurane vapors and inoculated in the left-rear footpad with a 10 μl volume containing 10^3^ PFU of virus diluted in PBS/1% FBS. Mice were weighed daily and disease scores were assigned as described previously (Jupille *et al*, 2011). In brief, the following criteria were used: score of 1: mild deficit in hind paw gripping of injected foot; score of 2: mild deficit in bilateral hind-paw gripping; score of 3: bilateral loss of gripping ability; score of 4: bilateral loss of griping ability, moderate bilateral hind-limb paresis, altered gait, difficulty righting self; score of 5: bilateral loss of gripping ability, severe bilateral hind-limb paresis, altered gait, inability to right self; score of 6: moribund state. At experiment termination, mice were euthanized by exposure to isoflurane vapors followed by bilateral thoracotomy. Blood was collected, mice were perfused with 5-10 mL of 1X PBS or 4% paraformaldehyde (PFA) (for experiments involving histology), and indicated tissues were harvested in *in vitro* diluent (1X PBS with 1% FBS and 1x Ca^2+^Mg^2+^) for analysis of infectious virus by focus formation assay (FFA) or plaque assay, or in TRIzol reagent Life Technologies) for RNA isolation and quantification of viral genomes by RT-qPCR. Tissues were homogenized using a MagNA Lyser instrument (Roche).

For serum clearance experiments, mice were anesthetized with isoflurane vapors and inoculated i.v. with 10^8^ genomes of CHIKV diluted in 100 μl of PBS/1% FBS. At 45 min post inoculation, mice were sacrificed and serum was collected. For lymph node accumulation experiments, mice were anesthetized with isoflurane vapors and inoculated s.c. in the left-rear footpad with a 10 μl volume containing 5 x 10^4^ PFU of virus. At 2 hpi, blood and the draining pLN were collected in TRIzol.

### Viral Genome Quantification by RT-qPCR

To quantify viral genomes, RNA was extracted from 20 μl of serum or from homogenized tissues in TRIzol reagent using the PureLink RNA mini kit (Life Technologies). CHIKV cDNA was generated from 10 μl of serum derived RNA or 1 μg of tissue derived RNA using random primers (Invitrogen) with SuperScript IV reverse transcriptase (Life Technologies). CHIKV genome copies were quantified by RT-qPCR using a CHIKV specific forward primer (5′-TTTGCGTGCCACTCTGG-3′) and reverse primer (5′-CGGGTCACCACAAAGTACAA-3′) with an internal TaqMan probe (5′-ACTTGCTTTGATCGCCTTGGTGAGA-3′), as previously described (Hawman *et al*, 2013). RRV cDNA was generated from 10 μl of serum derived RNA or 1 μg of tissue derived RNA using a sequence-tagged (indicated with lower case letters) RRV-specific RT primer (5*′*-ggcagtatcgtgaattcgatgcAACACTCCCGTCGACAACAGA-3*′*) with SuperScript IV reverse transcriptase (Life Technologies). RRV genomes were quantified by RT-qPCR using a tag sequence-specific reverse primer (5′-GGCAGTATCGTGAATTCGATGC-3′) with a RRV sequence-specific forward primer (5′-CCGTGGCGGGTATTATCAAT-3′) and an internal TaqMan probe (5′-ATTAAGAGTG TAGCCATCC-3′), as previously described (Stoermer *et al*, 2012).

### Plaque Assay and Focus Formation Assay

To quantify infectious virus, a plaque assay or focus formation assay (FFA) were used as previously described (Hawman *et al*., 2017). For plaque assays, samples were serially diluted 10-fold in 1X PBS + 2% FBS + 1X Ca^2+^Mg^2+^ and absorbed to BHK-21 cells in a 6-well plate for 1 h, after which cells were overlayed with 1% immunodifusion agarose (MP Biomedical). After incubation at 37°C for 40-44 h, cells were stained with neutral red stain and plaques were counted. For the FFA, serum or tissue homogenate were serially diluted 10-fold in 1X PBS+ 2% FBS+ 1X Ca^2+^Mg^2+^ and adsorbed to Vero cells in a 96-well plate for 2 h. Cells were then overlaid with 0.5% methylcellulose in MEM-alpha + 10% FBS and incubated at 37°C for 18 h. Following fixation with 1% PFA, cells were probed with CHK-11 monoclonal antibody (Pal *et al*, 2013) at 500 ng/ml diluted in Perm Wash (1x PBS, 0.1% saponin, 0.1% BSA), followed by a secondary goat anti-mouse IgG conjugated to horseradish peroxidase at 1:2,000 in Perm Wash. Foci were visualized with TrueBlue substrate (Fisher) and counted with a CTL Biospot analyzer using Biospot software (Cellular Technology).

### Immunohistochemistry

To evaluate KC depletion in the livers of DT-treated WT, Clec4F-DTR^+^, and CD169-DTR^+^ mice and PLL- or CLL-treated WT mice, at the time of harvest mice were perfused with 4% PFA and livers were harvested and fixed in 4% PFA for 24 h. Livers were paraffin-embedded and immunohistochemistry was performed on 5-micrometer sections using F4/80 antibody clone Cl:A3-1 (BioRad Cat. No. MCA497) and the VECTASTAIN Elite ABS HRP kit (Vector Laboratories, PK-6100) as previously described (Carpentier *et al*., 2019). To quantify the efficiency of KC depletion, F4/80^+^ cells were counted from 10 randomly selected high-power fields (HPF; 40X) for each stained liver section, and were used to calculated the average number of F4/80^+^ cells per HPF of view.

### Isolation of cells from lymph nodes and flow cytometry

To evaluate MARCO expressing cells in lymphoid tissue, popliteal and inguinal lymph nodes were pooled from WT or MARCO^-/-^ mice. Lymph nodes were minced with a 22-gage needle in 1 mL of digestion media (EHAA with 0.25 mg/mL Liberase DL and 17 μg/mL DNase) and incubated at 37°C for 1 h, after which an equal volume of dissociation buffer (0.1M EDTA in EHAA) was added and incubated at 37°C for 5 min. Cells were passed through a 100 μm cell strainer (BD Falcon). Single-cell suspensions were incubated for 20 min at 4°C with anti–mouse FcγRIII/II (2.4G2; BD Pharmingen) prior to staining for 1 h at 4°C with the following antibodies from BioLegend (most) or Novus Biologicals (MARCO) diluted in FACS buffer (PBS with 2% FBS): anti-CD45 (30-F11), anti-PDPN (8.1.1), anti-CD31 (390), anti-CD169 (3D6.112), anti-CD11c (BV510), anti-CD11b (M1.70), anti-B220 (RA3-6B2), anti-TCR*β* (H57-597), anti-F4/80 (BM8), anti-NK1.1 (PL136) and anti-MARCO (2359A). Cells were fixed overnight in 1× PBS/1% paraformaldehyde (PFA) and analyzed on a BD LSR Fortessa cytometer using FACSDiva software. Further analysis was performed using FlowJo software (Tree Star).

### Immunofluorescence and confocal microscopy

Lymph nodes were fixed in 1 mL of phosphate buffer containing 0.1 M L-lysine, 2% PFA, and 2.1 mg/mL NaIO4 at pH 7.4 for 24 h at 4°C, followed by incubation in 30% sucrose phosphate-buffered solution for 48 h, then in 30% sucrose/PBS for 24 hr. LNs were then embedded in optimal-cutting-temperature medium (Electron Microscopy Sciences) and frozen in dry-ice-cooled isopentane. Eighteen-μm sections were cut on a Leica cryostat (Leica Microsystems). Sections were blocked with 5% goat, donkey, bovine, rat or rabbit serum and then stained with one or more of the following: B220 (clone RA3-6B2, ThermoFisher), Lyve-1 (clone ALY7, ThermoFisher), CD169 (clone 3D6.112, BioLegend) MARCO (clone ED31, BioRad), CD11b (clone M1/70, BioLegend) and mCherry (polyclonal, Novus Bio, Cat# NBP2-25157). Images were acquired using identical photomultiplier tube (PMT) and laser power settings on a Leica Stellaris confocal microscope (Leica). Confocal microscopy images were collected over the entire popliteal lymph node (representing approximately a 7 mm^2^ imaged area) and individual fields (tiles) were merged into a single image file. Images were analyzed using Imaris v9.02 software (Oxford Instruments). Colocalization of Lyve1 and mCherry was performed using the Coloc module of Imaris (Oxford Instruments) and % of colocalized (double positive) voxels quantitated by the program using the same settings for each LN.

### Preparation of single-cell suspensions for single-cell mRNA sequencing

The draining popliteal lymph node from mock- or CHIKV-inoculated mice were pooled into individual replicates (3 replicates; LNs from 5 mice pooled per replicate). Lymph nodes were mechanically homogenized using a 22G needle in Click’s medium (Irvine Scientific, 9195) supplemented with 5 mg/mL liberase DL (Roche, 05401160001) and 2.5 mg/mL DNase (Roche 10104159001) for 1 h at 37°C. After incubation, digested tissues were clarified by passing through a 100 μm cell strainer. Cell suspensions were enriched for CD45^-^ cells by labeling cells with PE-conjugated anti-mouse CD45 (30-F11), CD140A (APA5), and Ter119 (Ter119) monoclonal antibodies and subsequent depletion of PE-labeled cells using Miltenyi anti-PE microbeads (130-048-801) and Miltenyi MACS LS (130-042-401) columns according to the manufacturer’s instructions with the following modifications: (1) we used 25% of the recommended volume of anti-PE microbeads and (2) we subjected the CD45^-^ enriched cell fraction to a second MACS LS column. All cell suspensions post-column enrichment were enumerated using a hemacytometer. Cell fractions throughout the procedure were analyzed for PE-labeled cell depletion and enrichment of CD45^-^ cells by flow cytometry. Cell fractions were stained with fixable LIVE/DEAD dye (Invitrogen, L34955) and antibodies against the following cell surface antigens: CD45 (30-F11), CD31 (390), PDPN (8.1.1), B220 (RA3-6B2), TCRβ (H57-597), CD11b (M1/70), and Ly6C (HK1.4). All flow cytometry antibodies from obtained from BioLegend, BD Bioscience or eBioscience. Following surface antigen staining, cells were washed, fixed in 1%PFA/1%FBS, and data was acquired on a BD LSR Fortessa X-20 flow cytometer. Data analysis was performed using FlowJo analysis software (Tree Star).

### Single-cell library preparation using the 10x Genomics platform

Lymph node cell suspensions enriched for CD45^-^ cells were subject to single-cell droplet-encapsulation using the Next GEM Chip G Kit (1000127) and a 10x Genomics chromium controller housed in our BSL3 laboratory. We targeted recovery of 10,000 cells for single-cell RNA sequencing for each replicate. Single-cell gene expression libraries were generated using the Next GEM single-cell 3′ GEM library and gel bead kit v3.1 (1000128) and single index kit T set A (1000213) according to the manufacturer’s instructions (10x Genomics). Sequences were generated with the Illumina NovaSEQ 6000 instrument using S4 flow cells and 300 cycle SBS reagents. We targeted 50,000 reads per cell, with sequencing parameters of Read 1:151 cycles; i7 index: 10 cycles; i5 index: 0 cycles; Read 2: 151 cycles.

### Transcriptome and oligonucleotide detection and analysis

FASTQ files for each replicate (3 mock, 3 CHIKV-infected) were processed using the cellranger count pipeline (v5.0.1). Reads were aligned to the mm10 and CHIKV AF15561 (EF452493.1) reference genomes. Analysis of gene expression data was performed using the Seurat R package (v4.0.0). Gene expression data for each biological replicate were combined into a single Seurat object. CHIKV counts were included as a separate “assay” in the Seurat object so they would not influence downstream processing (dimensionality reduction, clustering) of the mouse gene expression data.

CHIKV-low and -high cells were identified by first filtering cells to only include those with >5 CHIKV counts. K-means clustering was then used to independently group each biological replicate into CHIKV-low and -high populations (Walsh *et al*., 2021). Cells with 5 CHIKV counts or less were included in the CHIKV-low population. Cells were filtered based on the number of detected mouse genes (>250 and <6000) and the percent mitochondrial counts (<20%). Genes were filtered to only include those detected in >5 cells. Potential cell doublets were removed using the DoubletFinder (v2.0.3) R package using an estimated doublet rate of 10%. Due to the ability of CHIKV to inhibit host transcription (Fig. S8), CHIKV-high cells with a low number of detected mouse genes (<250) or a high fraction of mitochondrial reads (>20%) were not filtered and remained in the dataset for the downstream analysis. The fraction of CHIKV counts was calculated by dividing the number of CHIKV counts by the total number of counts for each cell.

Mouse gene expression counts were normalized by the total mouse counts for the cell, multiplied by a scale factor (10,000), and log-transformed (NormalizeData), Normalized mouse counts were scaled and centered (ScaleData) using the top 2000 variable features (FindVariableFeatures). The scaled data were used for PCA (RunPCA) and the first 40 principal components were used to find clusters (FindNeighbors, FindClusters) and calculate uniform manifold approximation and projection (UMAP) (RunUMAP).

B cells and T cells were identified based on CD19 and CD3 expression, respectively. The remaining cell types were annotated using the R package clustifyr (https://rnabioco.github.io/clustifyr) with published reference RNA-seq data (Heng, 2008; Malhotra *et al*., 2012; Rodda *et al*., 2018) available for download through the clustifyrdata R package, https://rnabioco.github.io/clustifyrdata). Endothelial cells were further classified using reference data for mouse LEC subsets (Xiang *et al*., 2020).

### Statistical Analysis

Appropriate experimental sample sizes were determined using a power calculation (80% power, 0.05 type I error) to detect a 4-5-fold effect in pre-existing sample sets. Each Figure legend defines the biological replicates of individual mice (N) and the number of experiments performed. Data are represented as mean ± SD or mean ± SEM, as indicated. The statistical tests conducted on each data set are indicated in the Figure legend and were performed using GraphPad Prism 8.0. Two-sided t-tests (parametric) or Mann Whitney tests (nonparametric) were used to compare two groups. One-way ANOVA with Bonferroni’s multiple comparison test (parametric) or Kruskal-Wallis with Dunn’s multiple comparisons test (nonparametric) were used to compare three or more groups, and two-way ANOVA with Bonferroni’s multiple comparison test was used to compare two groups at multiple time points.

## Acknowledgements

This work was supported by Public Health Service grants R01 AI148144 and R01 AI141436 to T.E.M., F32 AI140567 to K.S.C., F32 AI122463 to M.K.M., T32 AI007405 to K.S.C., T32 AI074491 to R.M.S, and R35 GM119550 and R01 AG071467 to J.R.H.. H.D.H. is supported by the Intramural Research Program of NIAID, NIH (https://www.niaid.nih.gov/) and R.M.S is supported by the RNA Bioscience Initiative. The funders had no role in study design, data collection and analysis, decision to publish, or preparation of the manuscript.

## Author Contributions

K.S.C., E.D.L., B.A.J.T., J.R.H., H.D.H., and T.E.M. designed the experiments. K.S.C., B.J.D., M.K.M., N.A.M, C.J.L, F.S.L., and G.V.R. performed the experiments. K.S.C., R.M.S., M.K.M., G.V.R., H.D.H., and T.E.M. performed the data analysis. K.S.C., R.M.S., and T.E.M. wrote the initial draft of the manuscript, with the other authors providing comments and edits to the final version.

## Declaration of Interests

The authors declare no competing interests

## Supplemental Figure Legends

**Figure S1. CHIKV tissue burdens in distal tissues are enhanced in MARCO^-/-^ mice at days 3, 7 and 14 pi. (A-C)** WT or MARCO^-/-^ C57BL/6 mice were inoculated subcutaneously in the left rear footpad with 10^3^ PFU of CHIKV. Viral tissue burdens were analyzed at 3 **(A)**, 7 **(B)** and 14 **(C)** days post inoculation (dpi) by FFA (A), or RT-qPCR (B and C). Mean ± SEM Two experiments for each time point, n= 10. Mann-Whitney test; \**P* < 0.05, \*\*\**P* < 0.001, \*\*\*\**P* < 0.0001.

**Figure S2. MARCO is expressed by medullary sinus macrophages and LECs in LNs.** LNs were pooled from uninfected WT or MARCO^-/-^ C57BL/6 mice. Representative flow plots and percentages of MARCO expressing cells by resident macrophage populations (A, C) and stromal cell populations (B, D) are shown. Mean ± SEM. Two experiments, n=4-5. Two-way ANOVA with Bonferroni’s multiple comparisons test; \*\**P* < 0.01, \*\*\*\**P* < 0.0001.

**Figure S3. Depletion of phagocytic cells in the lymph node and liver does not enhance CHIKV dissemination**. WT C57BL/6 mice were i.v. injected with PLL or CLL 42 h prior to virus inoculation and subcutaneously injected in the left rear footpad (FP) with PLL or CLL 24 h prior to virus inoculation as indicated. Mice were then inoculated s.c. with 10^3^ PFU of CHIKV in the left rear footpad, and tissues and serum were collected at 24 hpi. Infectious virus was quantified by FFA. Mean ± SEM. Two experiment, n=8. Two-way ANOVA with Bonferroni’s multiple comparison test, comparing all groups to IV PLL+ FP PLL group; \*\**P* < 0.01, \*\*\*\**P* < 0.0001.

**Figure S4. CHIKV-E2 mCherry is susceptible to clearance by a poly(I) sensitive scavenger receptor.** WT C57BL/6 mice were treated i.v. with poly(C) or poly(I) 5 min prior to i.v. inoculation of 10^8^ particles of CHIKV-E2 mCherry. Genomes in the inoculum and serum at 45 min post-inoculation were quantified by RT-qPCR. Mean ± SD. N=3, one experiment.

**Figure S5. CD45^-^ cell enrichment.** Pre- and Post-CD45^+^ cell depletion cell populations were analyzed by flow cytometry. (**A**) Representative plots of gating strategy used. (**B-C**) Representative flow plots of cell viability **(B**) and CD45^+^ and CD45^-^ cell populations (**C**) in pre- and post depleted postulations. **(D**) Percentages of CD45^+^ and CD45^-^ subsets among replicates. One experiment, n=3.

**Figure S6. LEC Annotations.** To assess the accuracy of endothelial cell type annotations, the subtype assignments were compared back to the reference data. The correlation with the reference RNA-seq data is shown for each subtype. Correlation coefficients (Spearman) are shown for each subtype.

**Figure S7. CHIKV-high classification and gene expression among CHIKV-high and CHIKV low cells. (A)** To identify cells with high amounts of viral RNA, cells were first filtered to only include those with >5 CHIKV counts. K-means clustering was then used to independently group each biological replicate into CHIKV-low and -high populations. Cells with <=5 CHIKV counts are included in the CHIKV-low group. CHIKV counts are shown below for each sample before filtering low quality cells (this includes all captured cells). **(B)** Cell quality metrics are shown for CHIKV-low and CHIKV-high cells for each replicate. These plots include all captured cells before quality filtering. CHIKV-high cells have fewer expressed mouse genes and an increased percentage of mitochondrial counts.

## Notes

### Competing Interest Statement

The authors have declared no competing interest.

