## Supplemental Figures for "MARCO^+^ lymphatic endothelial cells sequester arboviruses to limit viremia and viral dissemination"

**Figure S1**

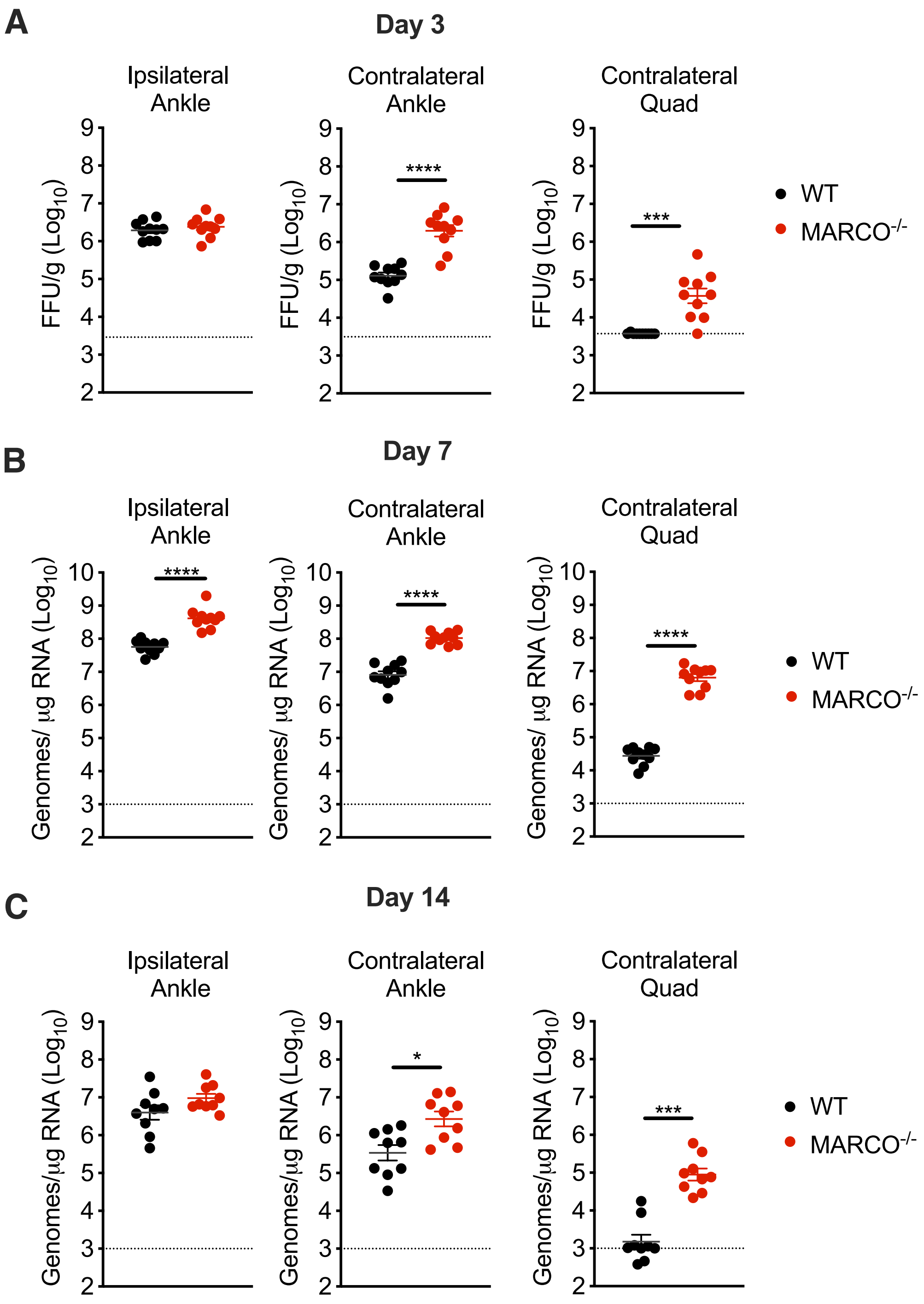


**Figure S1. CHIKV tissue burdens in distal tissues are enhanced in MARCO$^{-/-}$ mice at days 3, 7 and 14 pi. (A-C)** WT or MARCO$^{-/-}$ C57BL/6 mice were inoculated subcutaneously in the left rear footpad with $10^3$ PFU of CHIKV. Viral tissue burdens were analyzed at 3 **(A)**, 7 **(B)** and 14 **(C)** days post inoculation (dpi) by FFA (A), or RT-qPCR (B and C). Mean ± SEM Two experiments for each time point, n= 10. Mann-Whitney test; **P* < 0.05, ****P* < 0.001, *****P* < 0.0001.

**Figure S2**

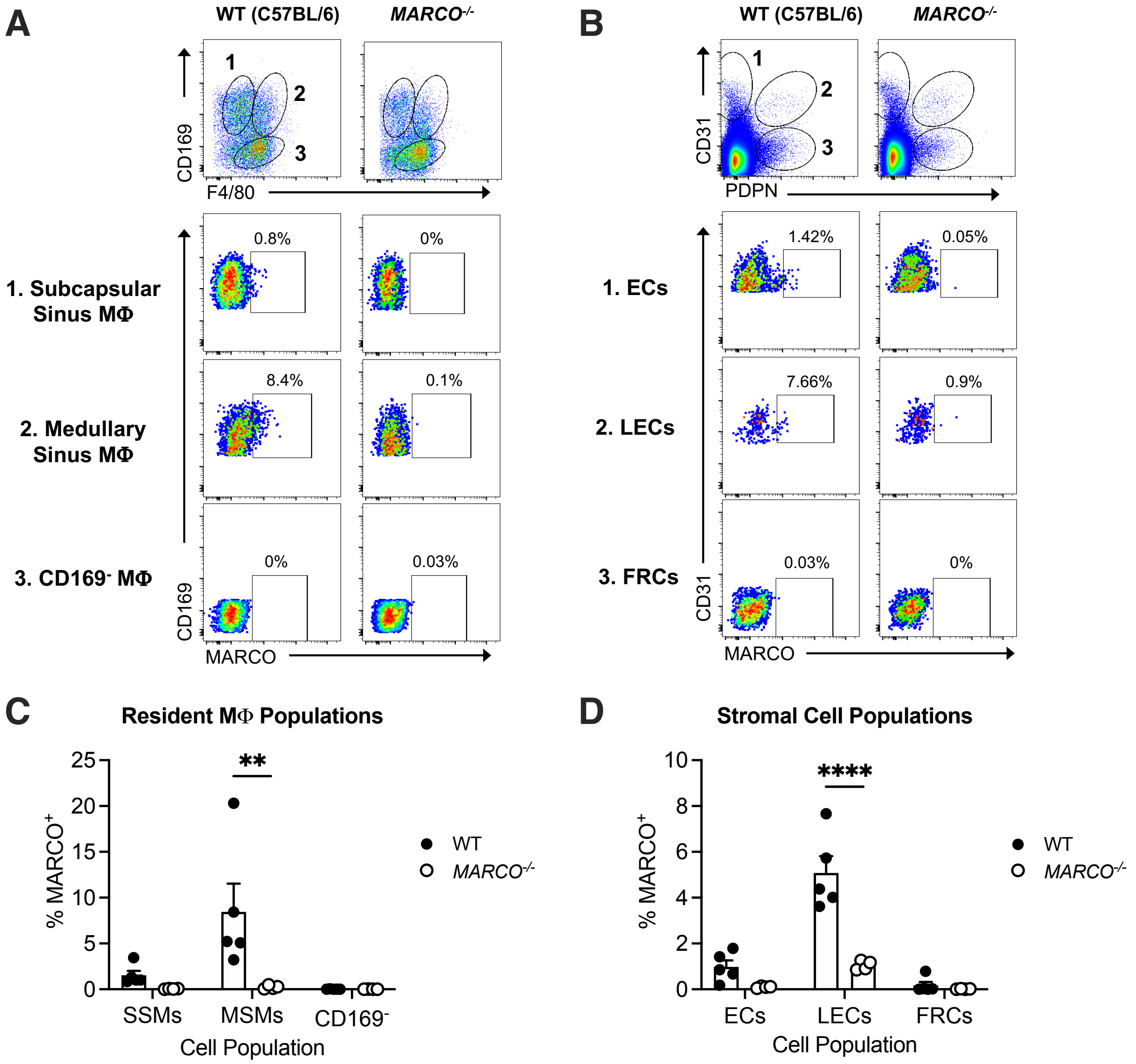


**Figure S2. MARCO is expressed by medullary sinus macrophages and LECs in LNs.** LNs were pooled from uninfected WT or MARCO$^{-/-}$ C57BL/6 mice. Representative flow plots and percentages of MARCO expressing cells by resident macrophage populations (A, C) and stromal cell populations (B, D) are shown. Mean ± SEM. Two experiments, n=4-5. Two-way ANOVA with Bonferroni's multiple comparisons test; $**P < 0.01$, $****P < 0.0001$.

**Figure S3**

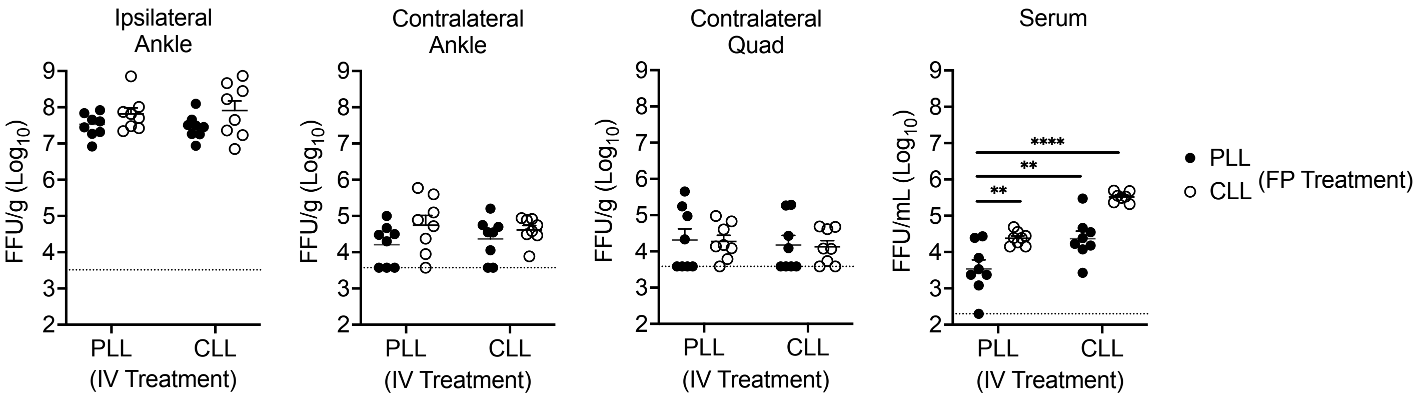


**Figure S3. Depletion of phagocytic cells in the lymph node and liver does not enhance CHIKV dissemination**. WT C57BL/6 mice were i.v. injected with PLL or CLL 42 h prior to virus inoculation and subcutaneously injected in the left rear footpad (FP) with PLL or CLL 24 h prior to virus inoculation as indicated. Mice were then inoculated s.c. with $10^3$ PFU of CHIKV in the left rear footpad, and tissues and serum were collected at 24 hpi. Infectious virus was quantified by FFA. Mean ± SEM. Two experiment, n=8. Two-way ANOVA with Bonferroni's multiple comparison test, comparing all groups to IV PLL+ FP PLL group; $^{**}P < 0.01$, $^{****}P < 0.0001$.

**Figure S4**

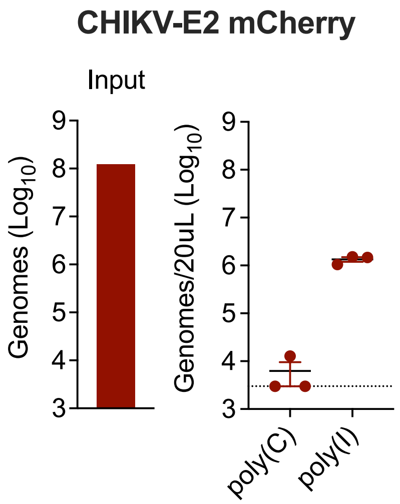


**Figure S4. CHIKV-E2 mCherry is susceptible to clearance by a poly(I) sensitive scavenger receptor.** WT C57BL/6 mice were treated i.v. with poly(C) or poly(I) 5 min prior to i.v. inoculation of $10^8$ particles of CHIKV-E2 mCherry. Genomes in the inoculum and serum at 45 min post-inoculation were quantified by RT-qPCR. Mean ± SD. N=3, one experiment.

### Figure S5

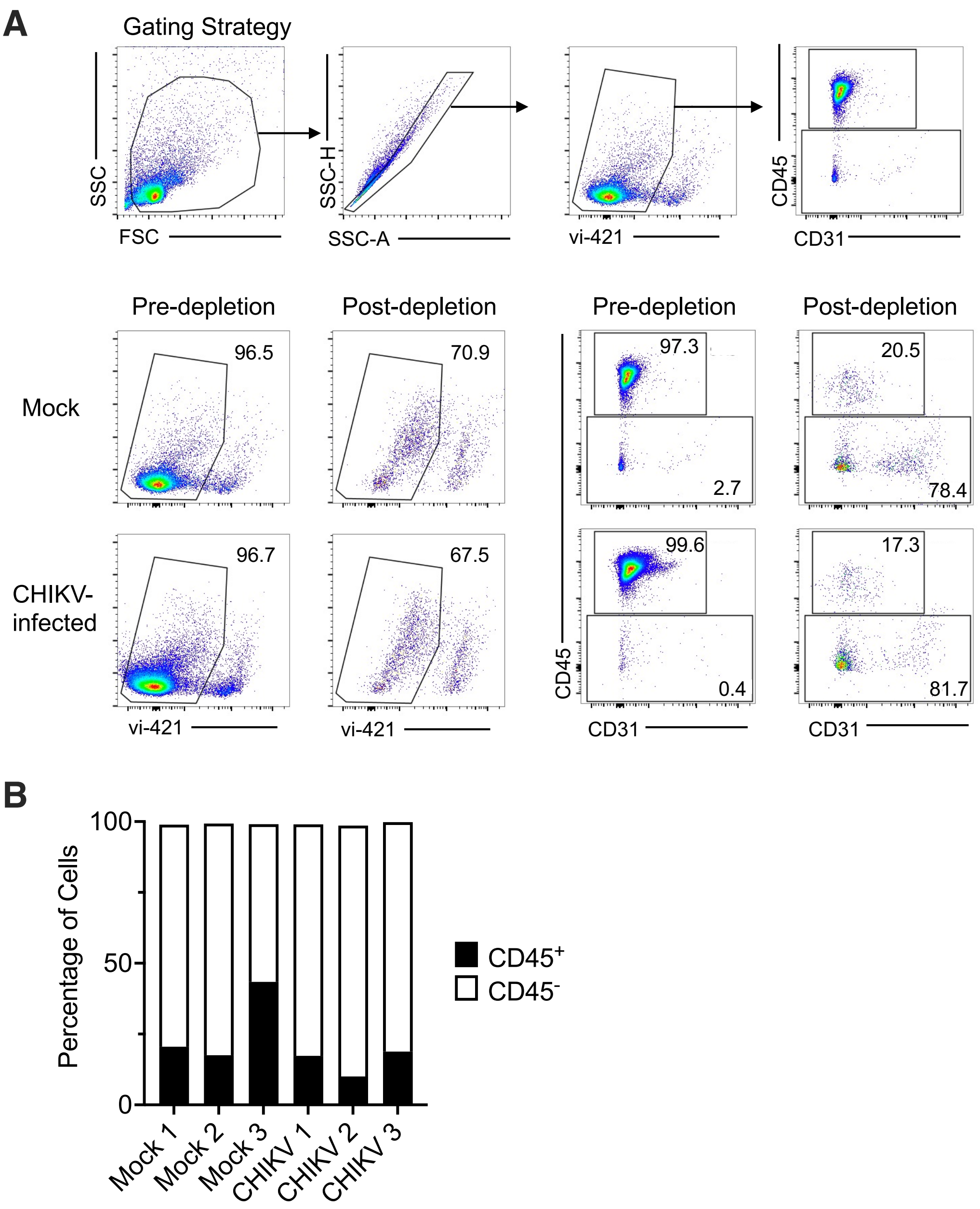


**Figure S5. CD45$^-$ cell enrichment.** Pre- and Post-CD45$^+$ cell depletion cell populations were analyzed by flow cytometry. (**A**) Representative plots of gating strategy used. (**B-C**) Representative flow plots of cell viability **(B**) and CD45$^+$ and CD45$^-$ cell populations (**C**) in pre- and post depleted postulations. **(D**) Percentages of CD45$^+$ and CD45$^-$ subsets among replicates. One experiment, n=3.

**Figure S6**

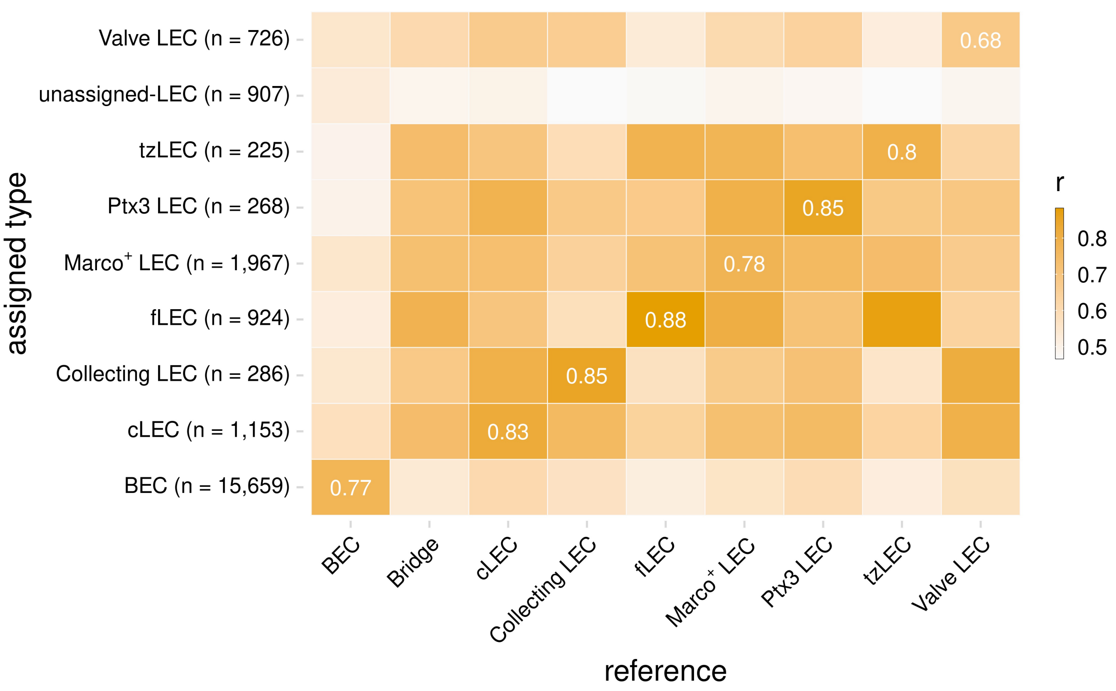


**Figure S6. LEC Annotations.** To assess the accuracy of endothelial cell type annotations, the subtype assignments were compared back to the reference data. The correlation with the reference RNA-seq data is shown for each subtype. Correlation coefficients (Spearman) are shown for each subtype.

### Figure S7

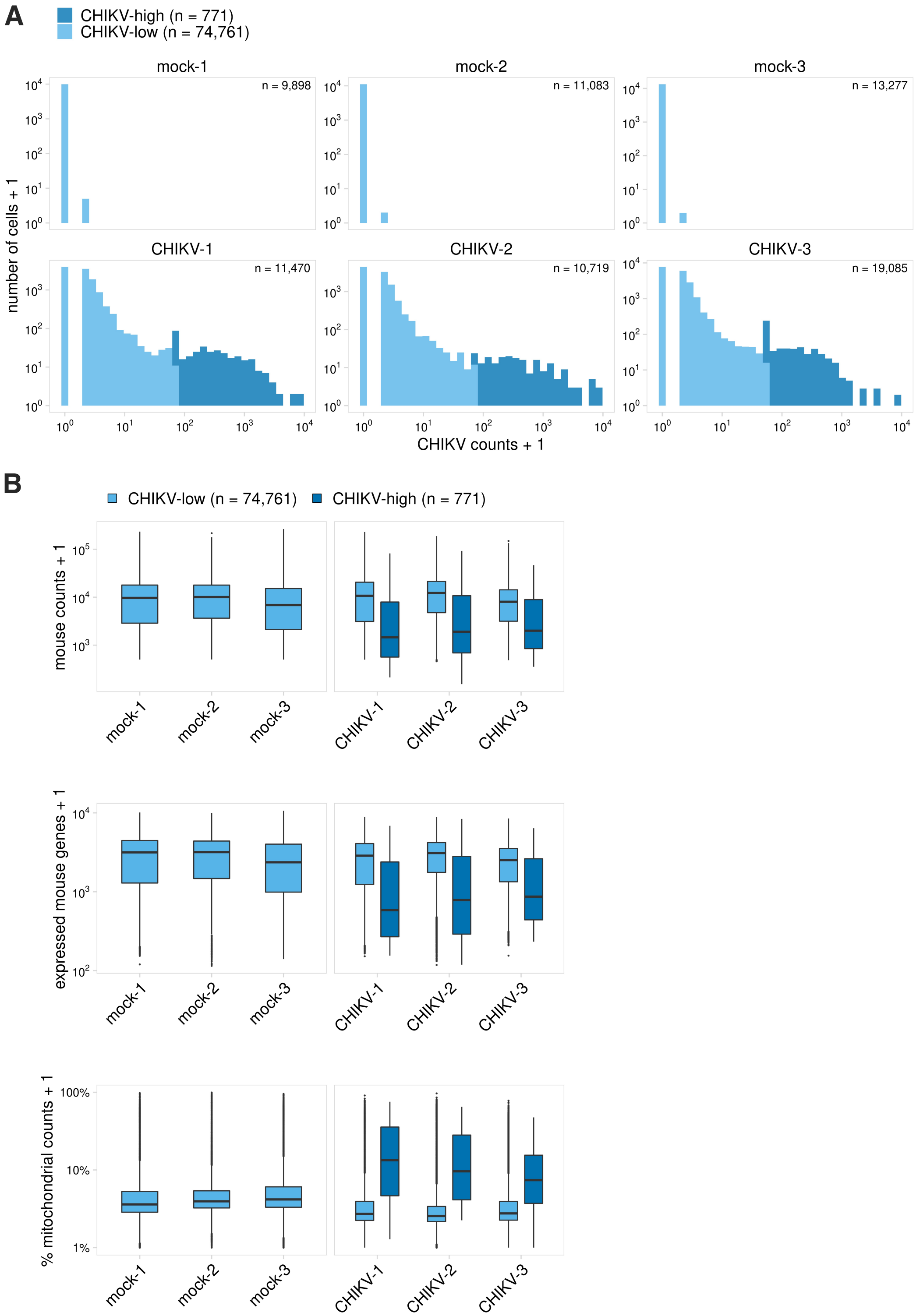


**Figure S7. CHIKV-high classification and gene expression among CHIKV-high and CHIKV low cells. (A)** To identify cells with high amounts of viral RNA, cells were first filtered to only include those with >5 CHIKV counts. K-means clustering was then used to independently group each biological replicate into CHIKV-low and -high populations. Cells with <=5 CHIKV counts are included in the CHIKV-low group. CHIKV counts are shown below for each sample before filtering low quality cells (this includes all captured cells). **(B)** Cell quality metrics are shown for CHIKV-low and CHIKV-high cells for each replicate. These plots include all captured cells before quality filtering. CHIKV-high cells have fewer expressed mouse genes and an increased percentage of mitochondrial counts.
